# Comprehensive multi-site profiling of the malignant pleural mesothelioma micro-environment identifies candidate molecular determinants of histopathologic type

**DOI:** 10.1101/2024.05.24.595690

**Authors:** David T Severson, Samuel Freyaldenhoven, Benjamin Wadowski, Travis Hughes, Yin P Hung, Lucien Chirieac, Roderick V Jensen, William G Richards, Ahmed A Sadek, Corinne E Gustafson, Kimberly Vermilya, Simona Inoccenti, Stephani Korle, Julianne S Barlow, Matthew B Couger, Jamie Anderson, Jason Meyerovitz, Vivian Wang, Mary N. Dao, Alex K. Shalek, Assunta De Rienzo, Raphael Bueno

**Author notes:** Corresponding author: Raphael Bueno, Division of Thoracic Surgery, Brigham and Women’s Hospital, 75 Francis Street, Boston, MA 02115, USA. These authors contributed equally to this work (David T Severson, Samuel Freyaldenhoven, Benjamin Wadowski).

## Abstract

Pleural mesothelioma (PM) comprises sarcomatoid, epithelioid and biphasic histologic subtypes. Bulk PM RNA-sequencing identifies a histology-associated molecular gradient with features of epithelial-mesenchymal (EM) transition but cannot parse malignant, stromal, and immune tumor components. The mechanisms driving PM malignant cell phenotype and associated histology is not well-characterized. Here, we use single-cell RNA-sequencing (scRNA-seq) paired with exome, bulk RNA-sequencing, and histologic analysis of adjacent samples to characterize malignant cell EM state, parse the tumor microenvironment (TME), and identify candidate drivers of PM cell fate. We observe EM variation in malignant cells analogous to bulk samples. We characterize epithelioid and sarcomatoid malignant cell programs and identify a new uncommitted malignant cell EM phenotype enriched in biphasic histology samples. Using inferred CNVs we observe that single individual PM clones consist of cells exhibiting all three EM cell states. We find that distinct non-malignant microenvironments associated with tumors consisting of mostly cells in each state, and identify *WNT* inhibition, *GAS6-AXL*, and *HBEGF-EGFR* signaling as pathways associated with distinct EM cell states. These findings provide deeper insight into the molecular drivers of PM malignant cells and identify non-malignant cell signals as potential EMT and growth drivers in PM.

## INTRODUCTION

Pleural mesothelioma (PM) is an incurable, near-universally lethal, and rare malignancy arising from the pleura. PM is associated with asbestos exposure (Carbone et al., 2019; Peto et al., 1982; Sekido, 2013). Asbestos has been banned in several countries, but its continued widespread use internationally and the decades-long latency period of PM pathogenesis mean PM incidence is unfortunately unlikely to decline soon(Alpert et al., 2019). PM may be divided broadly into three prognostic histologic types: epithelioid, biphasic, and sarcomatoid. Median survival ranges for each type are 12-27, 8 to 21, and 7 to 8 months, respectively (Yap et al., 2017). Additionally, overall survival benefit with immunotherapy treatment is greatest in non-epithelioid tumors (Baas et al., 2021; Nowak et al., 2022). Although histology provides insight into patient outcomes and response, the drivers behind histologic type are poorly understood. In particular, the mechanism behind biphasic cases, tumors with a mixture of the other histologic type, remains unknown.

Numerous studies in the last two decades have utilized next generation sequencing of bulk tissue samples to understand the heterogeneity of genetic and molecular perturbations across PM tumors (Severson et al., 2020). Collectively, these studies have built upon previous work to thoroughly characterize the predominant genetic lesions and identify epithelial-mesenchymal transition (EMT) and inflammation signatures as major axes of molecular variation in the disease (Alcala et al., 2019; Blum et al., 2019; Bueno et al., 2016; Hmeljak et al., 2018; Reyniès et al., 2014). Earlier work identified discrete molecular groups, but more recently the benefits of viewing PM tumors on a histopathologic epithelial mesenchymal (EM) gradient have been demonstrated (Alcala et al., 2019; Blum et al., 2019). A recent multi-omics approach also identified ploidy and methylation as additional important discriminating features (Mangiante et al., 2023).

Despite this extensive work characterizing PM, therapeutic options and predictive biomarkers remain unacceptably limited. Among several possible reasons frustrating translational efforts is the presence of substantial within-patient tumor and associated microenvironment heterogeneity. Another study explicitly examining key features in PM from two samples taken from a single tumor demonstrated distinct histomolecular and methylation phenotypes within an individual patient (Meiller et al., 2021). Although two samples may not provide a comprehensive description of *intra*-patient heterogeneity, clearly the PM tumor microenvironment (TME) is not constant even within a single patient’s tumor. Additionally, bulk studies are unable to parse which molecular features are driven by bona fide tumor cells or those found in the microenvironment. For this reason, the variation of malignant and stroma cells across and within patients remain poorly understood in PM. Specifically, the effect, if any, of genetic features and TME on tumor histopathologic state and, conversely, malignant cell specific features that sculpt the stroma and immune micro-environment are not well-described. Thus, a new map that parses the TME across multiple well-annotated tumor sites is necessary to navigate the complex and dynamic TME of PM and a critical step towards sorely needed efficient and successful drug discovery.

To that end, we performed in-depth examination of the TME of multiple anatomically mapped tissue samples freshly allocated from PM tumor resections using histopathology coupled with adjacent bulk and single-cell RNA-sequencing (scRNA-seq). To define truncal genetic lesions, we also obtained genomic profiles of each surgical case. With this approach, we demonstrate that previously reported histopathologic EM gradients are properties of bona fide malignant cells, define a new malignant cell specific EM gene program, and identify a gene program associated with malignant cell commitment to EM phenotype. We also identify variation in extra-cellular matrix (ECM) gene expression with associated clonal and immune micro-environment features.

## RESULTS

### Anatomically mapped multi-sample TME characterization of PM

We developed a workflow for fresh allocation of multiple anatomically mapped samples from surgical resections of PM tumors for in-depth TME characterization using scRNA-seq, bulk RNA-seq, and histopathologic analysis of immediately adjacent tissue (**Figure 1A**). In total, we collected 93 samples from 40 patients’ surgical resections including 3 negative-control pleura cases with non-PM pathology. We collected 2.3 samples on average from each case. The cases were distributed across anatomic site and diagnostic histology; the cohort included 19 epithelioid, 14 biphasic, and 3 sarcomatoid cases. One case was diagnosed on final clinical pathology after sample analysis as a peripheral nerve sheath tumor (PNS) instead of PM, and one of the 19 epithelioid case provided a post-treatment sample without viable tumor after allocation. These served as additional control data points in our analysis. Salient cohort features are described in **Table S1A**. For the 37 tumor cases, we performed parallel whole exome and optical genome mapping analysis to identify predominant single nucleotide (SNVs) and copy number variants (CNVs), respectively. We observed CNV and SNV lesions consistent with previous PM studies (**Figures 1B; Tables S1B-C)** (Bueno et al., 2016; Hmeljak et al., 2018; Jean et al., 2012; Mangiante et al., 2021). Histopathologic analysis of multiple sites revealed substantial variation in cellularity and morphology within and across tumors (**Figure 1C; Table S1E**). Adjacent bulk RNA-seq analysis confirmed our cohort was representative of previously observed EMT-related histomolecular variation (**Figure S9A-B**; **Table S1D** (Blum et al., 2019; Bueno et al., 2016; Severson et al., 2020).

**Figure 1:**
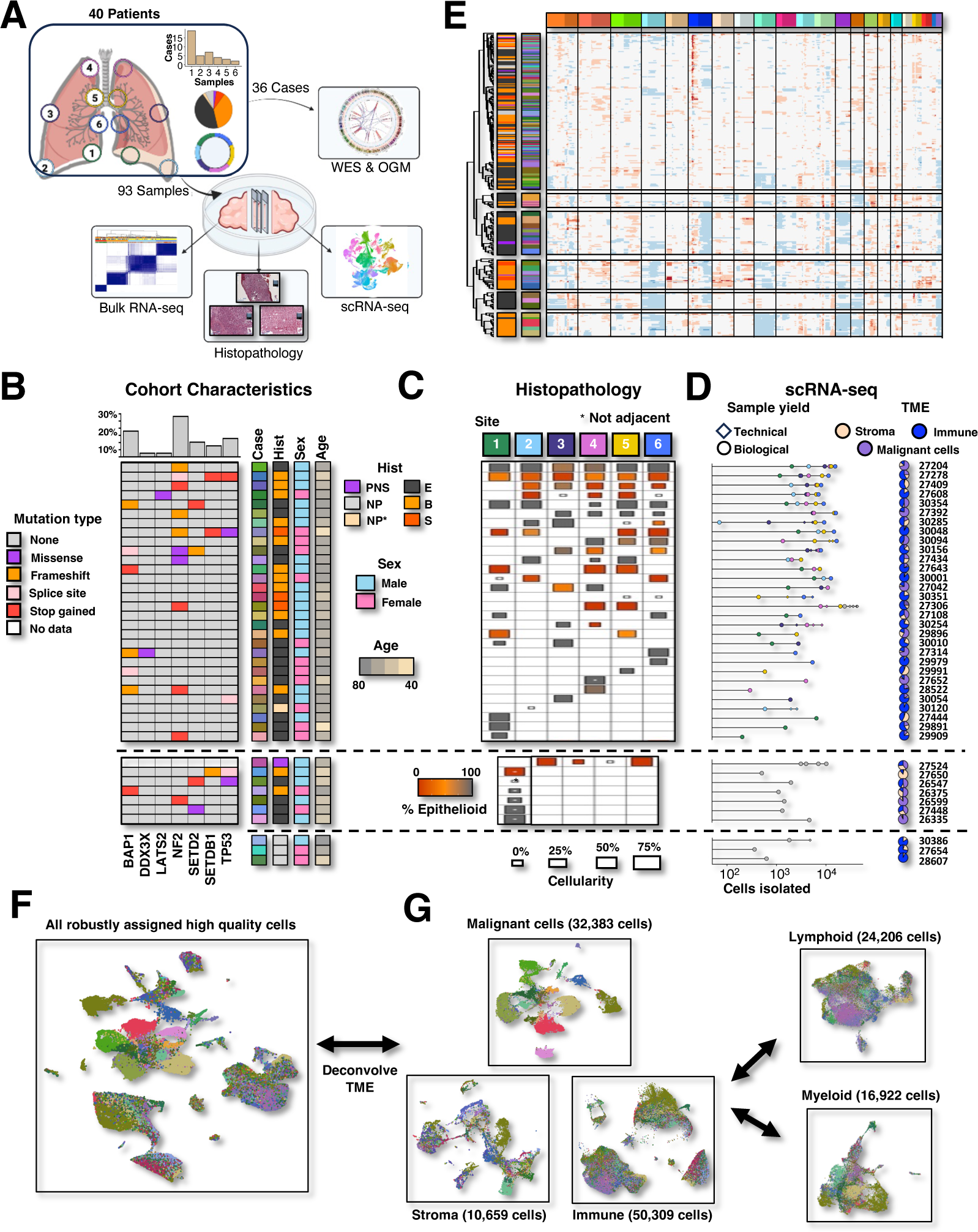
Comprehensive profiling of the MPM TME using scRNA-seq and multi-site sampling. **A)** Schematic illustrates experimental design for characterizing genomic profile of 40 cases and tumor micro-environment using bulk RNA-seq, scRNA-seq, and histopathology for 94 samples. The distribution of samples collected per case and the cohort histology are also displayed. **B)** Somatic mutations in known MPM driver genes are displayed in heatmap. Histology, sex, age, neoadjuvant therapy, average CV score, mean % epitheliod, and predominant transcriptional cluster are annotated by color. The barplot above summarizes the genetic perturbations to commonly mutated genes within the cohort and the line plot to the right displays the total SNV and CNV burden of samples. **C**) Pathologic analysis of samples for each case is displayed within each row, where slide size indicates cellularity and color indicates % epithelioid. An asterisk indicates that the examined slide is not immediately adjacent to scRNA-sequencing sample. **D**) Lollipop plot displays the number of cells isolated (length) in each case, where shapes denote cells isolated by biologic sample (large circle) or technical replicate (small diamond). Shape fill indicates anatomic site of sample. Pie charts display the TME distribution observed across cases. **E**) CNV structure of MPM clones as inferred by inferCNV and mapped to bands. **F)** UMAP plots display case of origin (color, 2B) of highest quality stroma (left) and tumor (right) cells with UMI>750 and genes>400.

We generated libraries for an average of 2,863 single cells per sample and 6,657 single cells per case (n=266,262 total cells; UMIs ≥ 300 & features ≥ 80) (**Figures 1D & S1; Table S1F**). The number of single cells isolated from each case and sample are depicted in **Figure 1D**. Dimensionality reduction and clustering coupled with marker gene analysis identified diverse immune and non-immune phenotypes (**Figure S2A-B; Table S2A; Methods)**. We observed both patient-ubiquitous and patient-specific clusters, where the latter implied the presence of non-malignant and malignant cells consistent with previously observed in human malignancy studies (**Figure S2C-D**) (Galen et al., 2019; Kim et al., 2018; Patel et al., 2014; Raghavan et al., 2021; Tirosh, Izar, et al., 2016). PM tumor cases were highly variable in their malignant, stromal, and immune cell content (pie charts in **Figure 1D**).

We thoroughly investigated these malignant cell calls by inferring CNVs from the scRNA-seq profiles as previously described **(Figures S3-4**) (Patel et al., 2014; Tirosh, Venteicher, et al., 2016). While broad phenotyping was readily accomplished with initial single cell library quality thresholds, stricter parameters (n=136,910 total cells with UMIs ≥ 750 and features ≥ 400) were required for inferring CNVs. After characterizing single cell CNVs, cells with CNV profiles suspicious for malignancy were re-analyzed and assigned to clonal groups as a function of their CNV profile (**Figures 1E**, **S5**). By combining cell phenotype, membership in patient-specific clusters, inferred CNVs, and clonal assignment, we were able to robustly assign malignant cells with explicit clonal assignment and infer malignant identity in the larger, less stringent cohort (**Figure S2 E-G**; **Methods**). Using the inferred malignant cell calls to assign microenvironment components, we observed wide ranging malignant, immune, and stromal cell content across PM tumors (**Figure S2F-G**).

### Malignant cells expression phenotypes encapsulate bulk EM gradient

We used the 32,383 high quality malignant cells (excluding the cases diagnosed as PNS and no malignancy) to characterize malignant cell specific expression of molecular pathways in PM (**Figure 1F**). Unbiased principal component analysis (PCA) revealed the EM gradient, ECM composition, inflammation, and cell-cycle to be primary axes of variation in PM malignant cells consistent with previous bulk studies (**Figures 2A-B & S6A)**. Next, we examined previously described bulk histomolecular signatures in this population of PM malignant cells (**Figure 2C**). We observed that bulk epithelioid- and sarcomatoid-related gene signature scores were highly concordant with the scRNA-seq signature. Histomolecular scores reflected diagnostic histology, slide histology, and previously assigned cell expression class; however, cells isolated from sarcomatoid cases more consistently separated from biphasic and epithelioid. Additionally, single cell expression of bulk EM gradient signature correlated strongly with analogous signatures in proximal bulk RNA-seq (**Figure 2C & S6A**). Taken together, these findings suggest that previously observed variation in bulk samples had significant contributions from PM malignant cell expression relative to stromal components.

**Figure 2:**
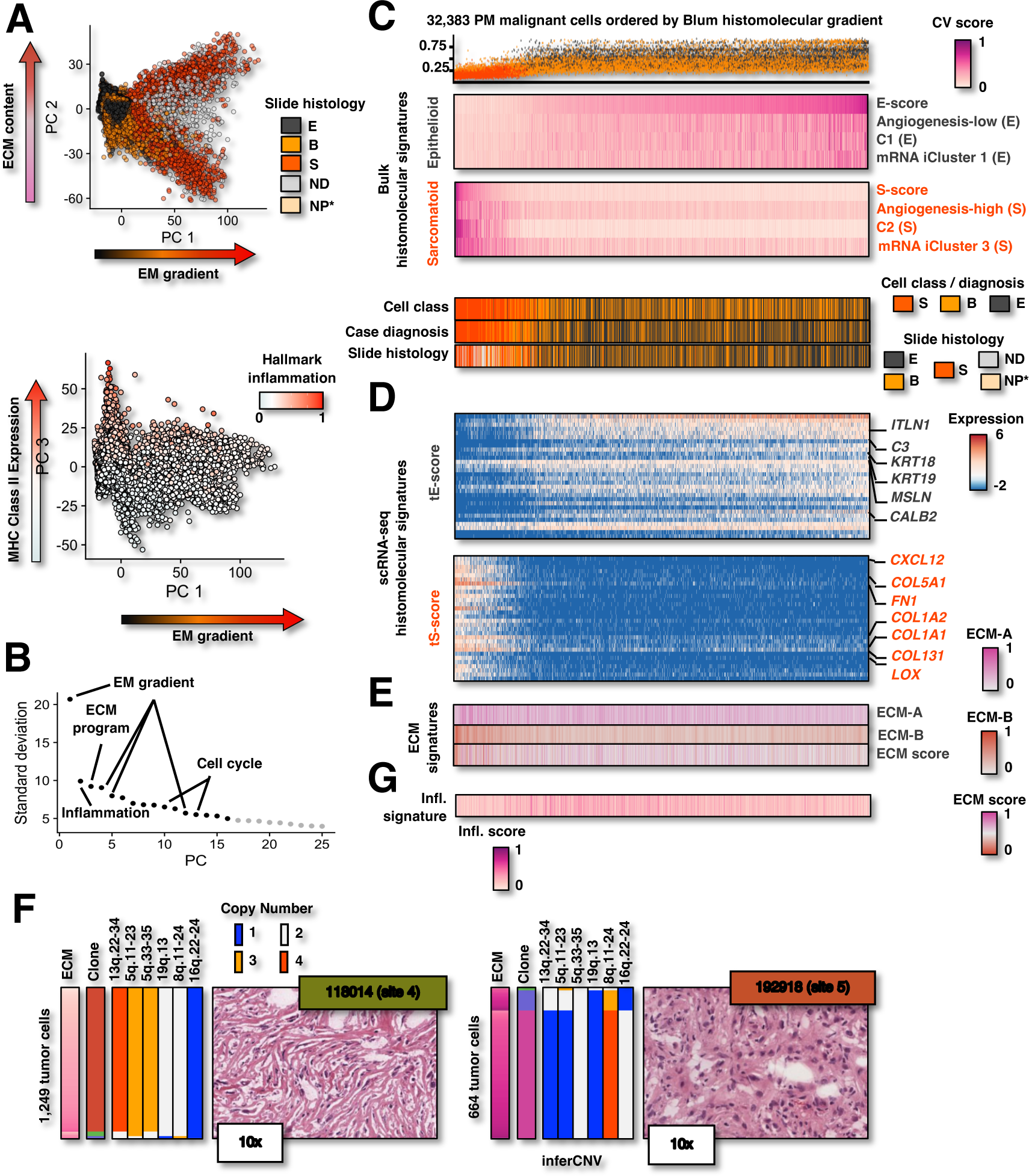
Malignant cells express previously described bulk EM gradient and distinct ECM programs. **A)** PCA of 32,383 tumor cells. Cells are annotated by immediately adjacent slide histology (top) or Hallmark inflammation score (bottom). **B)** Variation explained by principal component in malignant cell PCA analysis. **C)** Bulk histomolecular signature scores and cell phenotype features for 32.282 high quality inferred CNV confirmed MPM tumor cells ordered by the difference of Blum E- and S-scores. Cells are annotated by initial malignant cell transcriptional classification, PM case diagnosis, and adjacent slide histology. **D)** Cell-wise gene expression of top 30 tE- and tS-score genes. **E)** Cell-wise extra-cellular matrix signature scores. **F)** Morphology and architecture of adjacent H&E frozen sections are annotated with ECM phenotype, inferred CNV clone, and inferred CNVs of scRNA-seq profiles of cells isolated from immediately adjacent tissue (left) as well as OGM confirmed CNVs for each sample (right). Cells are ordered by clone and ECM score. **G**) scRNA-seq cell-wise scores for inflammation signature developed with principal component 3.

Because the EMT-related histomolecular gradient previously described in bulk samples (Blum et al., 2019; Severson et al., 2020) correlated significantly with primary axis of variation in bona fide PM malignant cells using unbiased analysis (r: −0.826, p-value < 0.001) (**Figure S6B**), we characterized a malignant cell-specific EM histomolecular gradient using the first principal component (PC) of these isolated malignant cell transcriptomes. That is, we identified genes that significantly correlated (|r| > 0.15) with a high (232 genes) or low (371 genes) first PC score to form malignant cell derived S- and E-score signatures (henceforth, tS-score and tE-score), respectively (**Figure 2D & S6C; Table S3D)**. Interestingly, only 20 (46.5%) and 28 (63.6%) of expressed Blum bulk signature S-score and E-score genes were present in these malignant cell-specific EM signatures. By isolating our analysis to malignant cells, we identified malignant cell-specific gene programs previously unreported in bulk studies illustrating the utility of a single-cell approach. KEGG pathways enriched in tS-score included “ECM receptor interaction”, “regulation of actin skeleton”, and “TGF-**β** signaling” while “antigen processing and presentation”, “complement and coagulation cascade”, and “glycolysis / gluconeogenesis” pathways were enriched in tE-score genes (**Table S3F**).

### Mesenchymal malignant cells express distinct ECM signatures

We also observed orthogonal variation (second PC) present in all samples but was especially evident in different tumor regions representing distinct CNV clones from a single sarcomatoid surgical case, 27306 (**Figures 2A, S5, S7A-B**). Excluding 27306 from PCA produced similar results suggesting that this variation was not driven by this sarcomatoid case alone (**Figure S7C**). To parse out EM gradient and case specific effects, we constructed shared and 27306-specific signatures from genes differentially expressed between cells at the extremes of PC 2 (**Table S4**). We annotated these signatures ECM-A (low PC 2) and ECM-B (high PC 2) to reflect the finding that numerous differentially expressed genes at either extreme were associated with the ECM, e.g., decorin/*DCN*, cathepsin B/*CTSB*, and lumican/*LUM* in ECM-A, and integrin beta 1/*ITGB1* and type VIII collagen A1/*COL8A1* in ECM-B (**Figure S7D-E**; **Table S4A**). This annotation was corroborated by unbiased pathway analysis that identified enrichment in focal adhesion, ECM-receptor response, and tight junctions in either signature (**Table S4B,E**). Most malignant cells had middling PC 2 scores and exhibited neither phenotype; these cells tended to be isolated from epithelioid tumors (**Figures 2E & S7F**). Intriguingly, in the 27306 sarcomatoid case, immediately adjacent histopathology revealed distinct architecture between two samples with different clone composition and ECM phenotype (**Figures 2F & S7I**). While ECM phenotype in this case clearly associated with cell clonotype (max clone difference: 0.679; p-value<0.001), the ECM phenotype of each clone also varied by sample (**Figure S7I**). This suggested that sample ECM, and thus architectural, phenotype was determined by both clonal composition and influence of microenvironment due to either the malignant component itself or infiltrating non-malignant cell perturbation of the milieu. Overall, these findings suggest that distinct ECM modules exist in PM independent of previously described EM variation, especially in sarcomatoid tumors, and these modules may influence tumor morphology and architecture.

The ECM-A and ECM-B modules, especially within case 27306, were also associated with specific chromosomal abnormalities and tumor immune microenvironment phenotype. Distinct clones isolated from anatomically disparate sites of 27306 expressed each module, which allowed contrasting of inferred CNVs to generate candidate causal CNVs. For ECM-A, these included losses of 13q, 5q, and 19q arms and 8q amplification. In contrast, the ECM-B phenotype was associated with 13q, 5q and 19q amplification and 16q loss (**Figures 2F & S5**; **Table S4C**). We observed similar CNV lesions in cells with similar ECM phenotypes isolated from other cases (**Figure S7G**). The tumor immune microenvironment also differed between these two sites. The 27306 sample with high ECM-B expression had a lower histologic inflammatory score and fewer *CD274*/PD-L1 expressing malignant cells (**Figure S7J; Table S1E**). In cells from all cases, ECM-A phenotype significantly correlated with both MHC class I (r=0.259; p<0.001) and II (r=0.493; p<0.001) expression (**Figure S7H**).

### PM malignant cells are variably committed to EM gradient cell fate

We observed a subset of malignant cells that equivalently expressed the E and S signature programs at low levels. These cells were not outliers in terms of UMIs detected, genes expressed, or mitochondrial transcript content and were detected in 27 of 28 cases (96.4%) with at least 50 high quality malignant cells per case (**Figures 3A-B & S8A**). This population had middling PC 1 scores and small differences in E- and S-scores suggesting that these cells were not committed to either phenotype (**Figure 3C**). Therefore, we computed the distance of all malignant cells from equal expression of E and S signatures, identified the ‘uncommitted’ malignant cells with this phenotype, and generated a histomolecular uncommitted signature consisting of genes associated with this cell population (**Figures 3D-E**). These uncommitted cells tended to be isolated from biphasic tumors (72.7% of cells) and to be actively cycling (18.2% of cells) compared to committed epithelioid (41.4% from biphasic, 6.8% cycling) and sarcomatoid type (7.1% from biphasic, 8.0% cycling) (**Figure S8B-C**). As seen previously, sarcomatoid cells exhibited a bimodal distribution of ECM phenotype (ECM-A vs ECM-B), but the epithelioid and this uncommitted cell population had middling ECM scores suggesting that the previously described ECM variation is largely a property of committed mesenchymal cells (**Figure S8D**). Finally, this cell population was enriched for a DNA methyltransferase (*DNMT3A*), a hydroxymethyltransferase (*TET1*), mesoderm specific transcript (*MEST*), and a sonic hedgehog ligand, *HHIP* (**Figure S8E**).

**Figure 3:**
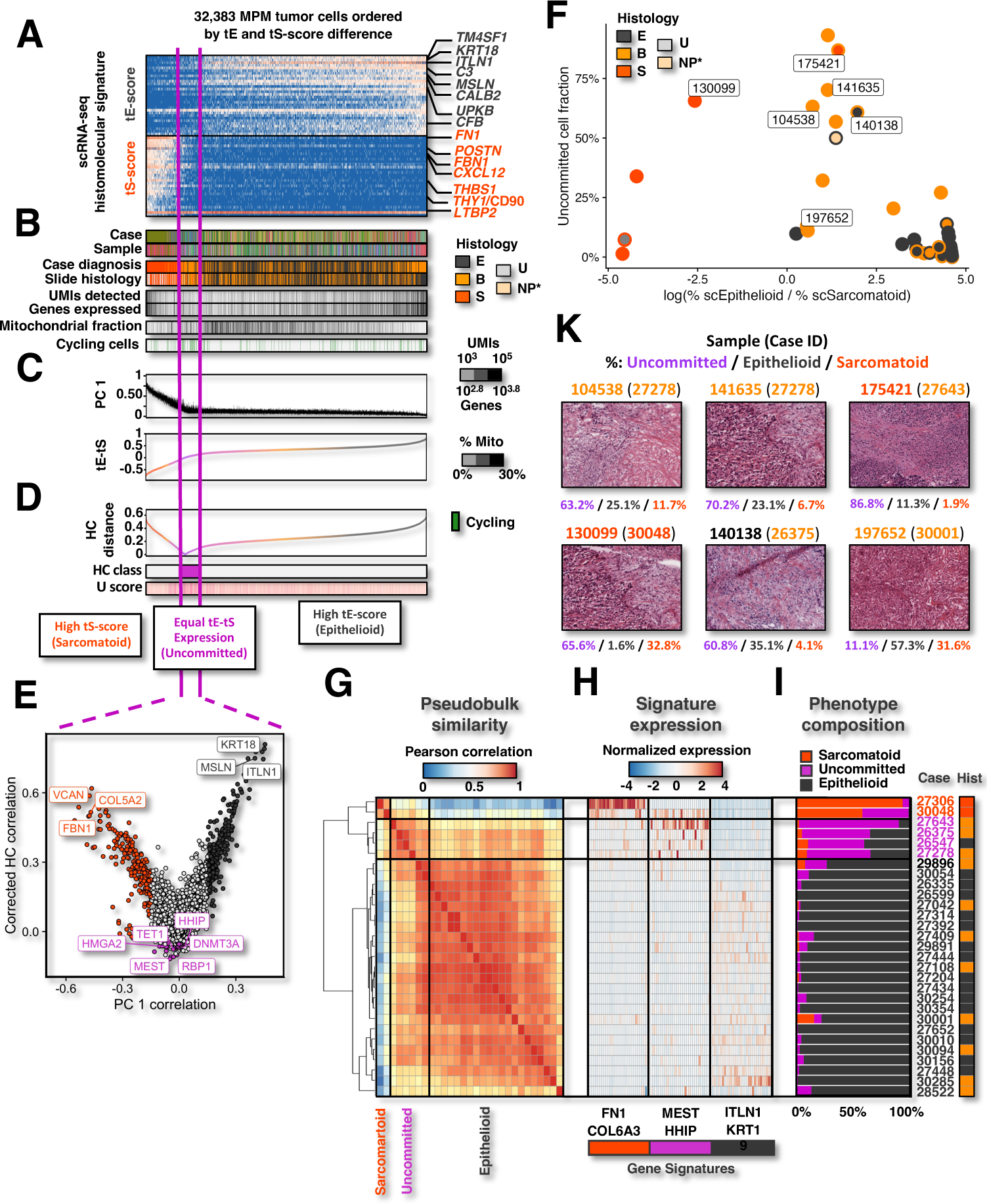
MPM malignant cells are variably committed to EM cell fate. **A)** scRNA-seq histomolecular signature scores and cell phenotype features (**B**) for 32,383 high quality inferred CNV confirmed MPM tumor cells ordered by the difference of Blum E- and S-scores. **C)** Principal component (PC) 1 and scRNA-seq EM scores for individual malignant cells. **D)** Histomolecular commitment (HC) distance, classification, and uncommitted (U) scores for malignant cells. **E)** Relationship between gene expression correlation to malignant cell corrected HC distance and PC1. Genes colored by phenotype program signature. **F)** Relationship between uncommitted and log ratio of epithelioid to sarcomatoid scRNA-seq malignant cell phenotype fractions in PM samples. **G)** Hierarchically clustered correlation of mean aggregated expression of scRNA-seq malignant cell phenotype signatures for each PM case (rows and columns). **H)** Mean aggregated expression of scRNA-seq malignant cell type signature genes (columns) for each PM case (rows). **I)** Fraction of cells by scRNA-seq malignant cell phenotype within each PM case (rows). Cases are annotated by histologic diagnosis. **K)** Morphology and architecture of adjacent H&E frozen sections are annotated with ECM phenotype, inferred CNV clone, and inferred CNVs of scRNA-seq profiles of cells isolated from immediately adjacent tissue (left) as well as OGM confirmed CNVs for each sample (right). Cells are ordered by clone and ECM score.

We examined the composition of uncommitted, committed sarcomatoid, and committed epithelioid malignant cells populations in each PM sample with at least 50 isolated malignant cells. Samples with higher fractions of uncommitted cells had similarly sized epithelioid and sarcomatoid committed populations (**Figure 3F**). Conversely, samples that skewed heavily sarcomatoid or epithelioid had a lower uncommitted cell fraction. Samples with the highest fractions of uncommitted cells were collected from PM cases with a biphasic diagnosis (**Figure 3F & S8F**). When examining cases in aggregate, we observed that malignant cells from PM tumors clustered into three groups, or pseudobulk phenotypes, expressing the sarcomatoid, uncommitted, and epithelioid malignant cell programs, respectively (**Figure 3G-H**). These clusters directly corresponded to the fraction of malignant cells of each type expressing these programs suggesting that bulk signatures represent a composition of these three discrete phenotypes (**Figure 3I**). Interestingly, a similar morphologic phenotype was observed in several samples, especially those isolated from case 27278, which had a particularly high fraction of uncommitted cells (**Figure 3K**). In contrast, the biphasic sample 197652, isolated from case 30001, contained high fractions of committed sarcomatoid and epithelioid malignant cells with relatively few uncommitted cells and exhibited correspondingly well-differentiated biphasic histology (**Figures 3F,K**).

We examined how malignant cell phenotype composition as measured by scRNA-seq associated with tissue-level features such as histopathologic and bulk transcriptional phenotype. Epithelioid and sarcomatoid malignant cell scRNA-seq phenotype composition significantly correlated (R=0.787, p=2.81×10^-9^) with histopathologic estimations of sarcomatoid and epithelioid content of adjacent slides (**Figure S8G**). Moreover, molecular cluster (Bueno et al., 2016) assigned to adjacent bulk RNA-seq samples trended with histopathologic and scRNA-seq phenotype compositions (**Figure S9D**). Samples with high fractions of uncommitted cells were assigned into the more mesenchymal molecular clusters 3 and 4 (**Figure S9E**). Together, these findings illustrate the utility of previous bulk classifications and pathologic analysis in capturing ratios of malignant cell phenotype despite confounding stroma and immune TME components. However, augmenting these approaches with scRNA-seq revealed the presence of a subset of phenotypically uncommitted cells which had not been previously described.

### Diverse non-immune stromal cells form the PM TME

To dissect malignant and stromal cell components of the PM TME, we examined high quality non-immune, non-malignant cells concomitantly isolated with malignant and immune cells (**Figure 1F-G**). The 10,659 high-quality isolated stromal cells included 668 non-neoplastic mesothelial cells, 3,386 fibroblasts, 2,456 endothelial cells, 856 vascular smooth muscle cells (vSMCs), 415 alveolar cells, 198 adipocytes, and 95 pericytes (**Figure 1G & S10A-D**; **Table S5A-B**). One hundred and thirty-six unfiltered macrophage cells formed a single cluster. An additional 2,154 fibroblasts predominantly isolated from the sarcomatoid 27306 case expressed MHCII programs representing either doublet or antigen-presenting fibroblast (apFibroblast) cells (Kerdidani et al., 2022). Other patient-specific clusters included 24 erythrocytes, 37 reactive endothelial cells, and 258 cells suspicious for epithelioid PM malignant cells. In general, stromal cell types were isolated from samples and cases across each histologic diagnosis (**Figure S10E**) and sample microenvironment histology (**Figure S10F**), especially when accounting for patient specific subpopulations which may represent uncalled malignant cells.

Fibroblast populations isolated from the cohort were diverse. In addition to 2 clusters identified as apFibroblasts, 10 other fibroblast subpopulations were observed. All 10 of these fibroblast subpopulations contained cells from multiple patients within our cohort indicating these were not patient-specific phenotypes and thus unlikely to be unidentified malignant cells (**Figure S10A-B; Table S5A-B**). Two fibroblast subpopulations expressed *TGFBI*; these had the strongest thrombospondin/*THBS2* expression (labelled ‘THBS+ fibroblasts’). Notably, ‘apFibroblasts’ with class II expression and ‘cycling fibroblasts’ with robust cell-cycle expression (*MKI67*, *TOP2A*) also expressed middling *TGFBI* in contrast to all other fibroblasts, which expressed *THBS2* minimally. Seven fibroblast subpopulations expressed insulin growth factor-1/*IGF1* and osteoglycin/*OGN*. Four of these, co-expressed *WISP2*, *CXCL14*, and *CFD*, where 2 of these 4 *WISP2*+ fibroblasts expressed *PCOLCE2* and *APOD*. Overall, these clustering and marker gene analyses suggested that fibroblasts infiltrating PM tumors have diverse and heterogenous expression phenotypes.

### PM immune microenvironment varies across PM tumors

After sub-setting high-quality cells and excluding multiplet or heat shock program dominated clusters, a total of 50,309 immune cells were isolated from PM samples (**Figure 1F-G**). We identified 2,132 B cells, 636 mast cells, 24,206 T- and NK cells, and 16,922 monocyte, macrophage, and dendritic cells (DCs) using cluster and known cell type marker genes (**Figure S11A-C; Table S6A-B**). Next, we separately sub-clustered the T- and NK cell and monocyte, macrophage, and DC groups to identify more granular subtypes of each of these populations (**Figure S11D-E, Table S6C-D**). PCA readily segregated monocytes, DCs, and macrophages (**Figure S11F**). A subset of macrophages expressed *MARCO* with or without *VCAN* and *FN1*, consistent with recruited and tissue resident alveolar macrophages, respectively (**Figure S11G; Table S6A- C**). Among tumor associated macrophages (TAMs), the expression phenotypes varied as two broad subtypes expressing distinct programs associated with either *SEPP1* or *SPP1* (**Figure S11H**) as previously observed in other cancer TMEs (Raghavan et al., 2021). We also observed that subsets of macrophages expressed either antiviral programs exemplified by *IFIT1* expression or pro-inflammatory programs including *THBS1*, *CXCL8,* and *IL1B* (**Figure S11G-H**). This variation was reflected in the resulting clusters (**Figure S11A-B, D**). Intriguingly, expression of antiviral programs was associated with *AXL* (**Table S6C**), a receptor tyrosine kinase known to promote EMT and proliferation in malignant cells and polarizes macrophages toward an immunosuppressive phenotype(Engelsen et al., 2022).

For NK and T-cells, sub-clustering and marker gene analysis identified subpopulations of CD4+ T cells, CD8+ T cells, and NK cells (**Figure S11A-B,E; Table S6A-B**). CD8+ T cells were differentiated by variable expression of features of exhaustion (i.e., *SLAMF7*, *HAVRC2*, and *PDCD1*) and cytotoxicity (i.e., *GZMB* and *PRF1*). One cluster of CD8+ T cells which we annotated as interferon CD8+ T cells (IFNr CD8 T cell) expressed anti-viral genes such as *IFIT1* and *MX1*. Among CD4+ T cells, we isolated regulatory (Treg), naïve, activated, and Th1 CD4+ T cells. Tregs were identified by *FOXP3* expression. Naïve and activated T cells were identified by the absence or presence of *TNF* and *IL4R*, respectively. *INFG* and *EOMES* were marker genes for the Th1 CD4+ subpopulation. Possibly because of the shared feature of cytokine program expression, the Th1 CD4+ T cell cluster also contained cytotoxic CD8+ T cells. Notably macrophage/T-cell doublet populations were also found in both sub-clustering analyses; these have been previously reported and may be biologic. These were annotated appropriately and excluded from further analysis.

### ScRNA-seq deconvolves benign and malignant mesenchymal signatures

To characterize non-malignant TME contributions to bulk RNA-sequencing signatures, we used differential expression analysis (**Table S5C**) to generate stromal and immune signatures for cell types identified in the microenvironment of our cohort (**Figures 2A, S10, & S11; Table S5D**). We minimized inclusion of genes expressed by malignant cells in the resulting stromal signatures by including malignant cell lineages as controls in our analysis (**Methods**). This analysis also generated a list of genes expressed by non-malignant cells in the micro-environment (**Table S5E**). Published bulk RNA-seq signatures were then assessed for genes expressed predominantly by microenvironment cell types. We found that all signatures contained numerous genes expressed by stromal and immune cells, where on average 36.9% of epithelioid and 66.9% of sarcomatoid bulk signatures contained genes associated with non-malignant cells (**Figure 4A & S12A, Table S5F**).

**Figure 4:**
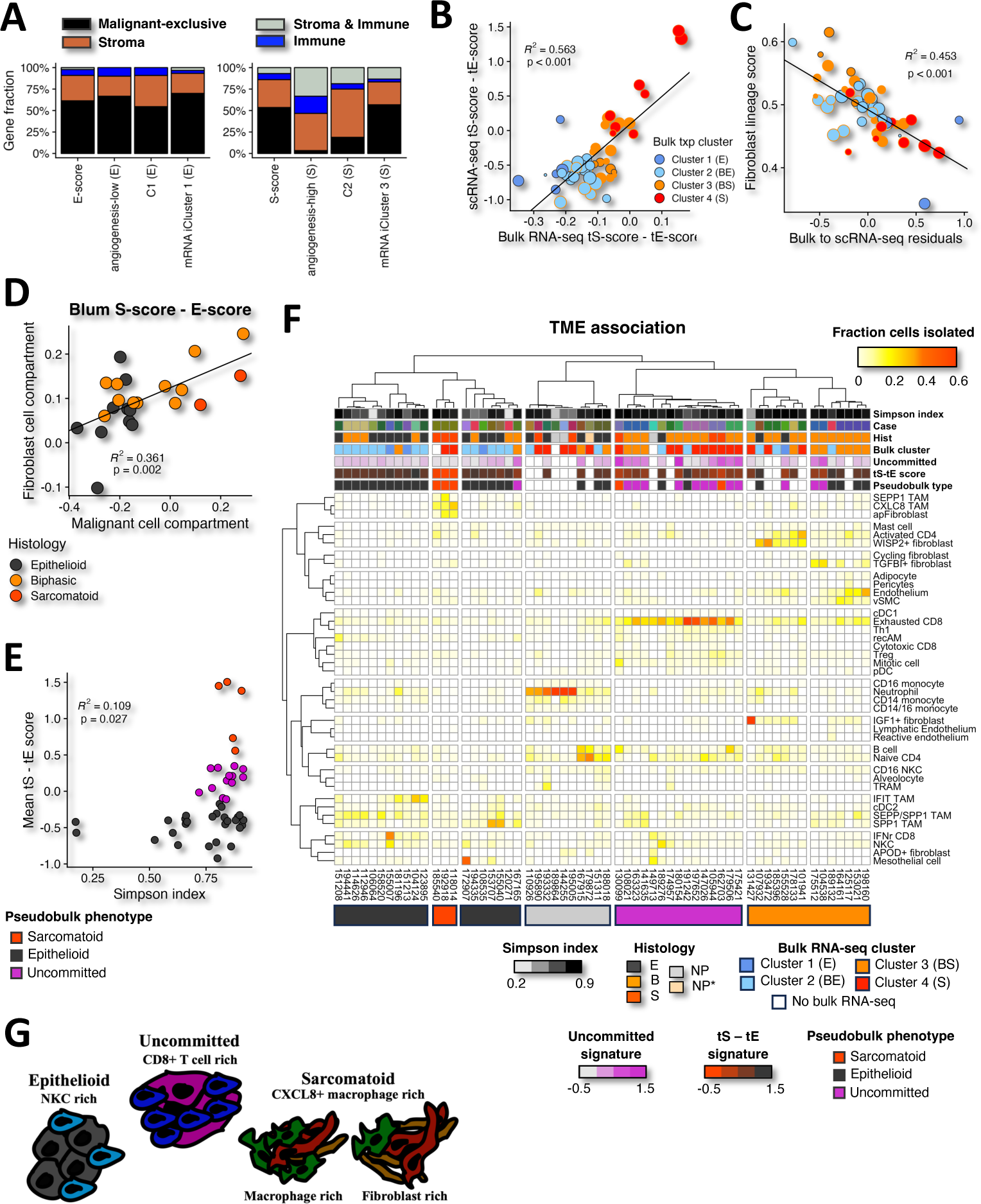
Non-malignant cells in the TME confound bulk signatures and associate with predominant malignant cell state. **A)** Barplots display cell compartments expressing published PM EM gradient gene signatures. **B)** Correlation between bulk scRNA-seq and scRNA-seq malignant cell EM gradient score estimates. **C)** Correlation between residuals of bulk to scRNA-seq EM gradient scores and fibroblast lineage score. Samples are annotated by bulk transcriptional cluster. **D)** Correlation between mean Blum S-score – E-score across fibroblast and malignant cells for each PM case with sufficient cells of each type. Case histology is annotated by color. **E**) Correlation between TME Simpson diversity index and scRNA-seq malignant cell signature. **F)** Heatmap displays fractional contribution of TME cell types, where samples and cell types are clustered and ordered by Pearson correlation. Samples are annotated with TME Simpson index, case of origin, case histology diagnosis (E: epithelioid, B: biphasic, S: sarcomatoid, NP:, NP*, bulk RNA-seq transcriptional cluster assignment, malignant cell uncommitted and tS-tE score signatures and pseudobulk malignant cell phenotype. Blank annotations indicated no malignant cells or adjacent sample existed to obtain that information. **G**) Model of TME associated with malignant cell state.

We assessed the impact of removing these genes from bulk EM cell state signatures. Specifically, we scored bulk transcriptomes of samples adjacent to scRNA-seq tissue using published gene signatures with and without genes expressed by non-malignant cells. Bulk signatures correlated significantly and strongly (p < 0.001 in all instances) with analogous signatures where genes expressed by non-malignant cells were removed (**Figure S12B-D**). This reinforces that malignant cell expression is the predominant signal underlying previous EM-gradient descriptions of bulk PM samples. Bulk sarcomatoid signatures performed universally worse than epithelioid signatures in terms of consistency; the average squared Pearson correlation coefficients were 0.895 and 0.988, respectively. This likely reflects the substantial overlap between bulk sarcomatoid signatures with non-malignant programs, especially fibroblast, and illustrates the importance of empiric micro-environment deconvolution by scRNA-seq. Additionally, bulk and scRNA-seq mean EM-gradient scores were concordant for the 51 scRNA-seq samples containing at least 10 malignant cells with bulk RNA-sequencing from the same sample site (**Figure 4B**, Pearson R^2^: 0.563; p-value<0.001). We observed that several samples had lower scRNA-seq tS-scores than would be predicted by their bulk score. These consisted mostly of previously described cluster 2 and 3 epithelioid and biphasic samples. We hypothesized that fibroblasts expressing some subset of genes within the sarcomatoid program at higher levels could explain this discrepancy. Indeed, we found that fibroblast lineage marker expression significantly correlated with the residuals computed in bulk and mean scRNA-seq histomolecular score linear regression (**Figure 4C**; Pearson R^2^: 0.453, p-value < 0.001). Thus, in a subset of PM tumors, benign mesenchymal expression modulates bulk tumor histomolecular gradient scores in addition to the predominant malignant cell signal. This further underscores the importance of explicit deconvolution of PM microenvironment cell type with scRNA-seq.

There is potential for diagnostic confusion between benign and malignant mesenchymal cells. Indeed, we observe that 58.2% of malignant tS-score signature genes (93.3% of the top 30 used for cell scoring) were among the 1,091 marker genes (adj. p-value < 0.001) for clusters identified as fibroblasts (**Figure S11E**). To clarify bulk molecular profile analysis and generate potential histologic biomarkers, we performed differential expression analysis between these two populations of single cells. We found that 140 fibroblast and 161 sarcomatoid genes were differentially expressed (FC >= 2 and adjusted p-value < 0.01) respectively (**Table S5F**). These included fibroblast markers of *C3*, *SPARCL1*, and *IGF1* as well as sarcomatoid markers of *CADM3*, *LTBP1*, and *CDH2* (**Figure S12F**). These gene sets can be used to deconvolve benign and malignant mesenchymal components of bulk PM microarray and RNA-sequencing samples for predictive and prognostic analysis. The genes also represent candidate biomarkers to distinguish benign fibroblasts and malignant sarcomatoid cells in PM samples.

In contrast, epithelioid malignant cells generally expressed mesothelial cell marker genes at higher levels rather than unique genes suggesting that epithelioid mesothelioma malignant cells express a similar gene program to non-neoplastic well-differentiated mesothelial cells (**Table S5F**). Genes differentially expressed in mesothelial cells were frequently mitochondrial genes. This is likely a reflection of the fragility of well differentiated mesothelium when compared to transformed epithelioid malignant cells.

Given the observed consistency between bulk and scRNA-seq scores and the large fraction of bulk sarcomatoid program genes expressed by fibroblasts, we hypothesized that EM-gradient expression in the non-malignant cells present in the TME mirrors malignant expression. Indeed, we observe that EM-gradient expression is significantly and strongly correlated between fibroblasts and malignant cells isolated from the same PM case (**Figure 4D**; Pearson R^2^: 0.361, p-value: 0.002) or tumor site (**Figure S12G;** Pearson R^2^: 0.376, p-value < 0.001). Significance was preserved when the malignant cell compartment was correlated with the entire stromal compartment (**Figure S12H**; Pearson R^2^: 0.401, p-value < 0.001), but not the immune compartment (**Figure S12I**; Pearson R^2^: 0.025, p-value: 0.200). These findings are consistent with the hypothesis that a paracrine factor within the TME drives PM mesenchymal tone for all tumor components.

### PM tumor microenvironment varies with sample histomolecular phenotype

Next, we assessed how PM features varied across multiple anatomic sites within a tumor. First, we assessed how malignant cell transcriptional state varies as a function of sample within cases. To accomplish this, samples were classified into epithelioid, sarcomatoid, and uncommitted pseudobulk phenotypes using correlation of aggregate expression of malignant cell EM state signatures (**Figure S13**). In general, the pseudobulk phenotype of samples isolated from the same patient were most strongly correlated with one another. Of the 15 patients with multiple samples, specimens from 7 of them more strongly correlated with other samples from the same patient rather than samples from other patients (**Figure S13E**). Only 4 cases consisted of samples assigned to distinct pseudobulk states. To classify the predominant malignant state in bulk samples, we developed malignant-exclusive signatures filtered by removing genes expressed by non-malignant cells in the TME (**see Methods**). The predominant malignant cell state of bulk samples was assigned by taking the state with the highest score in bulk samples. These bulk-assigned malignant states were mostly consistent (92.3% concordance) with pseudobulk malignant type of adjacent scRNA-seq tissue (**Figure S14A**). This suggests that most tumors have a predominant EM phenotype despite potentially large physical distances between isolated samples from single PM tumor. We similarly assigned 211 previously published bulk RNA sequencing profiles (Bueno et al., 2016), which demonstrated the presence of tumor samples predominated by each of the three malignant cell states in an orthogonal cohort (**Figure S14B**).

Distinct immune and histopathologic axes of variation have been proposed with multi-omics analysis (Mangiante et al., 2023). However, the interaction between immune and histopathologic tumor phenotypes has not been well-described in the setting of bulk sample analysis. To understand how sample microenvironment interacts with malignant cell state, we analyzed the non-malignant microenvironment. First, we assessed overall TME diversity by computing Simpson’s index for the 60 samples with at least 200 isolated non-malignant cells. Among 45 samples with sufficient non-malignant and malignant cells, we found that increasing diversity was significantly associated with more mesenchymal tumors (**Figure 4E**; R^2^: 0.109, p-value: 0.027). However, this result may be confounded by weak correlation observed between scRNA-seq EM signatures and the number of isolated cells (R^2^: 0.087, p-value: 0.042). Certainly, a subset of epithelioid tumors demonstrated less diversity than the rest of the cohort.

We then clarified the relationships between stromal and immune cell types by correlating relevant marker gene expression average over isolated cells of each type to understand relationships between cell types (**Figure S15A-B**). As expected, related cell types clustered together. APOD+ and, to a lesser extent, TGFBI+ fibroblasts were more unique in their expression profile than closely related IGF1+ and WISP2+ fibroblasts (**Figure S15A**). Cycling and apFibroblasts shared expression features with both endothelial/pericyte populations and other fibroblasts subtypes. For tumor-associated macrophage subtypes, SEPP1+ and CXLC8+ TAMs were distinct from the more closely related SEPP1/SPP1 co-expressing, SPP1+, and IFIT TAMs (**Figure S15B**).

After clarifying the relationships between TME cell types, we then clustered the 60 PM samples with greater than 200 non-malignant cells isolated (**Figure 4F**). Interestingly, clustering on TME composition segregated tumors into groups predominated by distinct pseudo-bulk malignant cell states. A group of low malignant cellularity samples containing two of the normal pleural controls contained higher fractions of WISP2+ fibroblasts, neutrophils and circulating monocytes. Explicit comparison of cell type fractions within the TME identified exhausted CD8 T cells to be significantly enriched in PM tumors predominated by uncommitted state (**Figure S15C**). Although not significant, higher fractions of TGFBI+ fibroblasts, naïve CD4, and SPP1+ TAM were also observed in tumors predominated by uncommitted malignant cell state. Committed epithelioid samples were significantly depleted in regulatory T cells (Treg), but significantly enriched with IFN receptor positive CD8 T cells (IFNr CD8) and natural killer cells (NKCs). In contrast, committed sarcomatoid samples were significantly enriched in CXCL8+ TAM.

ScRNA-seq data facilitates deconvolution of individual cell types but is costly, limiting the number of samples. Additionally, specific populations may be differentially isolated as a function of dissociation and processing time. Therefore, as aforementioned we developed cell type gene signatures for malignant cell, stromal, and immune cell types that allow bulk RNA-seq samples to be assessed for cell type composition (**Figure S14, Table S5C-F, Methods**). We scored bulk RNA-seq tissue samples adjacent to scRNA-seq samples with these cell type scores (**Figure S14C**). We observed that clustering on the TME composition segregated PM tumor samples by predominant malignant cell state suggesting that malignant cell state phenotype and the associated TME are related. Uncommitted samples were generally rich in fibroblasts and poor in macrophages. Additionally, these tumor samples were rich in CD8+ T cells, especially exhausted T cell phenotypes. Sarcomatoid and low cellularity samples were also enriched for macrophages, especially CXCL8+ macrophages. Interestingly, biphasic samples with high epithelioid content were rich in TGBI+ cells in adjacent bulk RNA-seq samples.

As with proximal bulk samples of the scRNA-seq samples, bulk malignant state was assigned using malignant exclusive scRNA-seq phenotype scores in 211 previously published bulk RNA-seq samples (Bueno et al., 2016) (**Figure S14B**). Next, we assessed scRNA-seq derived TME cell type scores in these 211 samples. While scRNA-seq scores of bulk RNA samples provided did not discriminate subtypes, i.e. cytotoxic vs exhausted CD8+ T cells, in the larger cohort of 211 bulk RNA-seq samples, broad cell type distributions were observed (**Figure S14D**). Malignant cell state again segregated by TME phenotype reinforcing the finding that malignant cell and associated TME phenotype are related in PM. Intriguingly, we observed that mesenchymal tumors were broadly divided into fibroblast rich, macrophage poor and fibroblast poor, macrophage rich groups. Explicit comparison revealed that in this orthogonal cohort with 7 sarcomatoid samples, CXCL8+ macrophages were indeed enriched in samples predominated by sarcomatoid malignant cell state compared to uncommitted (p-value: 0.0038) and epithelioid (p-value: 2.31×10^-6^) phenotypes (**Figure S15D**). Uncommitted-dominant samples were also enriched for exhausted CD8+ cells compared to epithelioid (**Figure S15E**; p-value: 5.68×10^-5^).

Thus, both scRNA-seq and bulk RNA-seq analysis in an independent cohort demonstrate that malignant cell state is associated with a distinct TME (**Figure 4G**). Specifically, sarcomatoid samples are enriched for CXCL8+ macrophages, where some sarcomatoid tumors vary in their overall fibroblast to macrophage ratio. Uncommitted-predominant samples are enriched for infiltration of CD8+ T cells, especially those expressing markers of exhaustion. Epithelioid tumors frequently have less TME cell type diversity and have an enrichment of natural killer cells.

### *MEST* and *HHIP* operationalize uncommitted cell state

Although the multi-gene meHCscore successfully assigns uncommitted cell state in bulk RNA sequencing data, we assessed top marker genes (**Figure S8C**), *MEST* and *HHIP*, to identify single gene biomarkers of this newly identified PM malignant phenotype. We observed that cases predominated by uncommitted cell state had significantly higher bulk RNA-seq MEST expression than epithelioid cases (**Figure S16A**, p-value: 0.0409). Notably MEST was sensitive, identifying all uncommitted rich cases, but not specific as it was also highly expressed in a subset of committed samples (Figure S16B). Interestingly, committed samples with high *MEST* expression had middling EM states as measured by Claudin-vimentin (CV) score (**Figure S16C**). Even in low uncommitted samples, MEST may be a biomarker for mixed malignant state PM tumors. The biomarker HHIP was similarly significantly enriched in uncommitted cases (**Figure S16D**; p-value: 0.006). Although HHIP was more specific and still denoted mixed state PM cases (**Figures S16E-F**). The biomarkers also identified uncommitted tumors in an orthogonal cohort of 211 bulk RNA-seq samples (**Figures S16G-H**). In these 211 samples, both *HHIP* and *MEST* were significantly elevated in cases with inferred uncommitted malignant cell state compared to epithelioid and sarcomatoid. Using a representative biphasic sample, we observed in spatial transcriptomic analysis that MEST demarcated a unique malignant cell population distinct from epithelioid and sarcomatoid malignant cells (**Figure S17**).

### Individual PM clones span the EM gradient

To assess how malignant cell state varied by clonotype, we examined the malignant cell EM score across individual clones within PM cases. We observed that while EM state was frequently restricted by case of origin, it generally did not vary by inferCNV defined clone within PM samples (**Figure 5A**). Indeed, cells from individual clones often expressed two or three malignant cell states. In one uncommitted biphasic case, five of six clones expressed all three malignant cell phenotypes (**Figures 5B**). In this case, clones were found to represent genuine subclonal heterogeneity as confirmed by orthogonal optical genome mapping (**Figure 5C**). Thus, EM malignant cell phenotype is not restricted by specific malignant cell clones and varies among cells that share the same copy number features. However, we did observe that EM malignant cell phenotype was distinct among clones in a handful of cases from each histology (sarcomatoid: 30048, biphasic: 26547, and epithelioid: 29891, 30010). Thus, while distinct genetic features are not a prerequisite for PM malignant cell heterogeneity, there may be specific lesions that restrict or modify EM malignant cell state.

**Figure 5:**
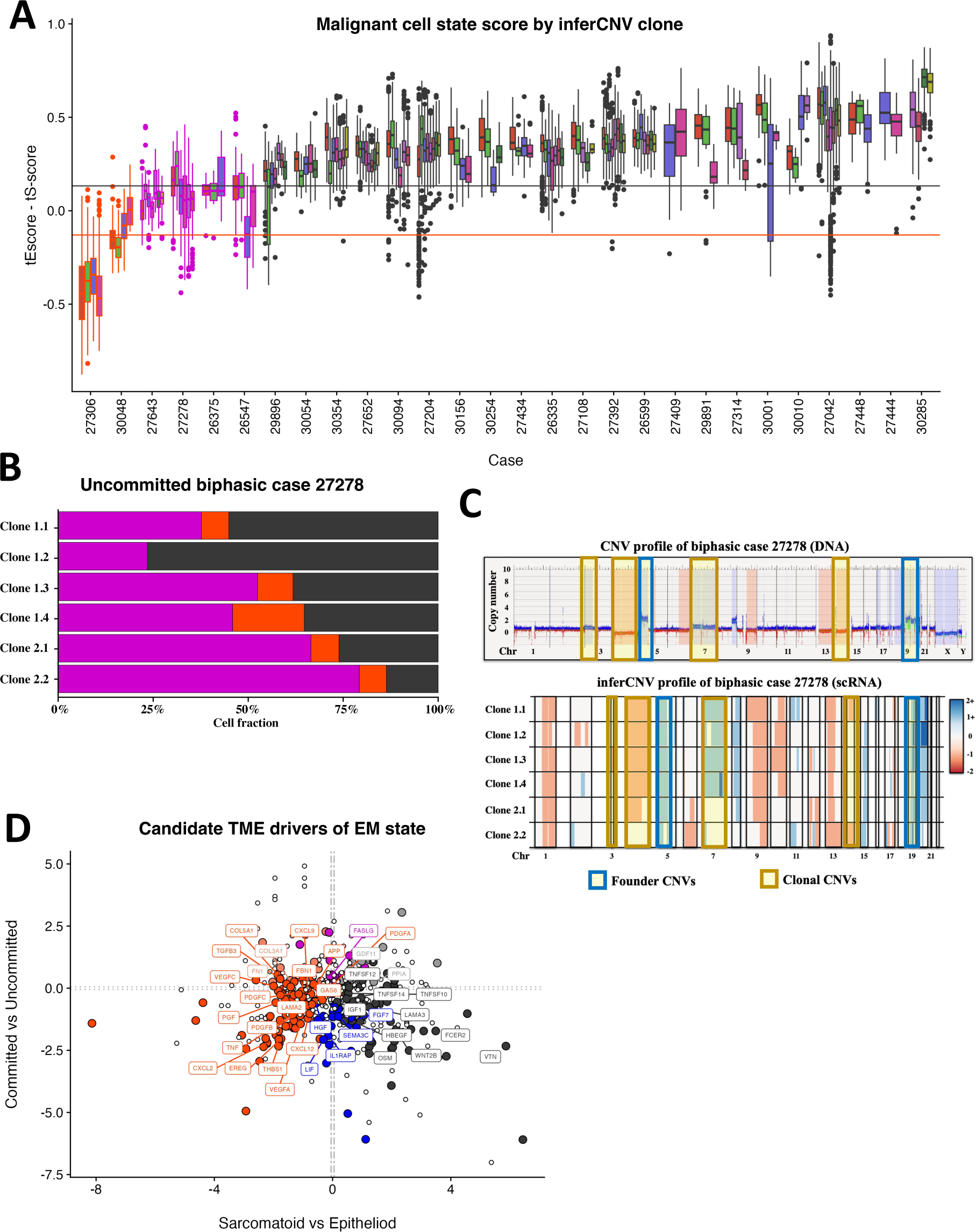
PM EM gradient represents a plastic phenotype with candidate TME drivers. **A)** Boxplot displays EM gradient score distribution of malignant cells across PM cases. Box outline and fill display scRNA-seq pseudobulk state and inferred CNV clone, respectively**. B**) Proportion of cells in each malignant cell state within six distinct inferred clones of biphasic case, 27278. **C**) Optical genome mapping (top) and inferred CNV profile of biphasic case 27278. **D)** Log2 fold change gene expression of signal proteins in non-malignant cells between tumors of distinct pseudobulk malignant cell states. Labelled genes were identified as having significant TME interactions using CellPhoneDB.

### TME factors associate with EM gradient

Because EM malignant cell phenotype varied within PM clone, we hypothesized that factors within the TME may drive EM cell state. To identify potential TME drivers, we performed differential expression analysis comparing extra-cellular signal expression between tumors predominated by each malignant cell phenotype. To parse extrinsic drivers expressed by non-malignant cells in the TME and intrinsic ones expressed by malignant cells themselves, we contrasted signal expression separately in non-malignant (**Figure 5D; Table S7A)** and malignant (**Figure S18A; Table S7B**) cells. We further narrowed candidate drivers with CellPhoneDB by filtering differentially expressed genes for those identified as components of significant interactions between malignant cells with either themselves or other cell types in the TME (**Table S7C**). We found that sarcomatoid malignant cell state was associated with TME expression of *CXCL2, CXCL12*, *VEGFA, THBS1, TGFB3*, and *GAS6* (**Table S7D**). *THBS1*, *CXCL12, VEGFA* and *GAS6* were also significantly more expressed by sarcomatoid malignant cells themselves along with *TGFB1* and *VEGFC* (**Table S7E**). *CXCL2* and *CXLC12* were generally expressed by myeloid cells, but mostly in CXCL8+ TAMs and alveolocytes, respectively (**Figure S18B**). These were identified as interacting with *DPP4* receptor expressed by malignant cells (**Table S7C**; **Figure S18C**). *VEGF* and *THBS* were mediated by *NRP1*/*NRP2* or integrin expression in malignant cells, respectively, and both widely expressed by sarcomatoid malignant cells, myeloid cells, and fibroblasts (**Figure S18B-C**). *TGFB3* interacted with the TGB1 receptor and was expressed by pericytes, vascular smooth muscle cells, *WISP2*+ and *TGFBI*+ fibroblasts. Finally, *GAS6* was expressed by sarcomatoid cells, dendritic cells, and SEPP1 TAMs. It was identified as interacting with the *AXL* receptor, which was highly expressed in sarcomatoid cells. These enriched signals provide new candidate biomarkers for sarcomatoid PM and candidates for drivers of PM mesenchymal cell state.

Epithelioid predominant tumors were notable for their *HBEGF* expression mediated by *EGFR* and expressed ubiquitously by myeloid cell types. Intriguingly, epithelioid malignant cells had significantly higher *WNT2B* expression. In contrast, uncommitted and sarcomatoid cells express the WNT inhibitory receptors *SFRP2* and *SFRP4*, respectively. This could represent a mechanism, whereby the more mesenchymal malignant states downmodulate beta-catenin. Interestingly, WNT pathway inhibitors *DKK1*, *DKK2*, and *DKK3* were also upregulated in uncommitted and to a lesser extent, sarcomatoid malignant cells (**Table S7B**).

## DISCUSSION

Using a large, histologically representative cohort of multi-site PM samples, we have demonstrated that previously described EM histomolecular variation in bulk PM samples exists within malignant cells themselves. To our knowledge, this is the first scRNA-seq PM cohort with matched bulk RNA-sequencing and adjacent histologic samples with a multiple-site sampling design. While previous studies in bulk RNA identified a continuous EM gradient across samples (Alcala et al., 2019; Blum et al., 2019), these analyses were unable to identify which cell components were driving this phenotype. The concordance between bulk and malignant cell-specific EM signatures demonstrated in the present study suggests that previous findings in bulk tissue were largely representative of malignant cell variation and is consistent with another recent PM scRNA-seq study (Giotti et al., 2024). Furthermore, we describe a new malignant cell state, termed uncommitted, in PM most frequently isolated in biphasic tumor samples. We identified malignant cell specific molecular signatures for EM, characterized by the expression of *ITLN1*, *KRT19*, and *C3* or *COL1A1*, *FN1*, and *LOX* at the epithelial or mesenchymal ends of the spectrum, respectively. Uncommitted cells expressed *MEST, HHIP*, *DNMT3A,* and *TET1*. Additionally, we describe candidate genetic determinants of ECM component variation in mesenchymal samples that associate with immune microenvironment. Given that tumor architecture has been associated with tumor staging in other malignancies, e.g., non-small cell lung carcinoma, these ECM modules may have further implications in prognosis as well.

We also demonstrate that individual malignant cell states are not defined as distinct clones. Instead, individual PM clones may generate all three phenotypes. This does not preclude genetic determinants of malignant cell state but does provide evidence that distinct mutations are not required for changes in PM cell state. Indeed, by untangling the PM micro-environment with scRNA-seq, we identify distinct drivers of proliferation and possibly EMT expressed by both malignant cells and their surrounding micro-environment. Specifically, we identify fibroblasts, pericytes, and myeloid lineage cells as cell types that express secreted drivers of EMT and proliferation. This is an intriguing finding given PM is a cancer of tumor suppressor mutation. While mutations remove the checkpoint on proliferation, it may be that non-malignant cells provide the fuel for proliferation, where mesenchymal cells tend to be driven by *GAS6-AXL* and epithelioid tumors by *HBEGF-EGFR* signaling. Notably, *GAS6-AXL* signaling has been previously identified as a an important EMT gene in PM (Engelsen et al., 2022; Ou et al., 2011)and *AXL* was recently described as associated with sarcomatoid malignant cell module gene expression in another PM scRNA-seq study(Giotti et al., 2024). We also identify inhibition of WNT as a potential mechanism for maintaining an uncommitted cell state, given the identification of *SFRP2, HHIP*, *DKK1*, *DKK2*, and *DKK3* as either signature biomarkers or significantly differentially expressed in in uncommitted cells. Given these findings, modulation of non-malignant signaling may provide a new drug target pathway in PM.

The single-cell RNA-sequencing approach utilized in this study is limited by the presence of substantial noise characteristic of such experiments. While every effort was made to remove multiplet cells, some libraries will inevitably contain expression from two or more distinct cells. Additionally, for many cells, expressed genes may not have been detected and these so-called “dropouts” may confound phenotypic and inferred CNV classification. The inferred CNVs provide a powerful window into malignant cell clone architecture, especially in PM where the CNV burden is high; however, these profiles are subject to the cell’s chromatin state and fluctuations in gene expression which may generate incorrect CNV calls. Parallel Optical Genome Mapping (OGM) does largely overcome this limitation by verifying genuine CNVs in many cases, but it could incorrectly eliminate true inferred CNVs that are clonal and not sampled in the tissue allocated to OGM. While the cohort was representative of all PM histologic types, the analysis was limited by the collection of only three sarcomatoid samples, where only two of these yielded viable malignant single cell libraries. Although bulk RNA-seq and IHC sections were taken from adjacent tissue, the tumor micro-environment may change even over a few microns of tissue. As a result, although this work was designed to minimize potential discordance between parallel modalities due to sampling, it may not have eliminated this effect entirely.

## ACKNOWLEDGEMENTS

This work was supported solely by the International Mesothelioma Program at Brigham and Women’s Hospital. The study sponsors played no role in the study design, collection, analysis, interpretation of data, writing of the report, or decision to submit the paper for publication.

## AUTHOR CONTRIBUTIONS

**Conceptualization:** D.T.S., S.F., R.B., W.G.R., A.DR.; **Validation:** A.A.S., W.G.R.; **Formal analysis:** D.T.S., R.V.J.; **Investigation:** D.T.S., S.F., B.W., Y.P.H., R.V.J.; **Resources:** T.H., A.DR., W.G.R., C.E.G., A.A.S., T.P.A, S.G., K.V., J.S.B, J.A., J.M., V.W., M.N.D., M.C.G.; **Data curation:** W.G.R., A.DR., C.E.G., A.A.S., D.T.S.; **Writing – original draft:** D.T.S.; **Writing – reviewing & editing:** D.T.S., R.V.J, R.B., W.G.R., Y.P.H., B.W., S.F. **Visualization:** D.T.S., B.W.; **Supervision:** R.B., A.K.S.; **Project administration:** R.B., D.T.S.; **Funding acquisition:** R.B.

## METHODS

### PATIENT CONSENT AND TISSUE COLLECTION

All samples were obtained from surgical resections at Brigham and Women’s Hospital from 2018-2021 under the Institutional Review Board (IRB) approval (Dana Farber Cancer Institute Protocol No 98-063). Patients provided written informed consent for allocation, banking, and downstream analytics and data publication. Using a common anatomic mapping scheme (**Figure 1A**), surgeons performed systematic multi-site pleural sampling at the time of macroscopic complete resection irrespective of gross malignant cell involvement at each site. In most cases, six-site sampling (the prespecified maximum) was achieved. Fresh samples were then processed by clinical pathology, and remaining tissue was allocated to the Brigham and Women’s Tissue and Blood Repository Core for research. At the core, the largest sample from each site was trisected. Half was allocated fresh for single cell transcriptomics, while the immediately adjacent tissue of each side was oriented in OCT and banked for bulk RNA-sequencing and histopathological analysis, respectively. Additional fresh frozen tissue was stored and annotated by the Institutional Tumor Bank with IRB approval (#98-063) at the Brigham and Women’s Hospital. Malignant cell content was confirmed by a pulmonary pathologist who reviewed H&E-stained frozen sections.

### BULK DNA AND RNA EXTRACTION

Bulk DNA for exome sequencing was isolated using the DNeasy Tissue kit (Qiagen, Hilden, Germany). Matched normal DNA was prepared from peripheral blood or lung tissue. For RNA extraction, the TRIzol (ThermoFisher Scientific, Waltham, MA) method in combination with RNeasy kit (Qiagen) was conducted. RNase or DNase I (Qiagen) treatments were conducted according to the manufacturer’s instructions. Nucleic acids were quantified using an NanoDrop One spectrophotometer (ThermoFisher Scientific). The integrity of the DNA was determined using Qubit 4.0 Fluorometer (ThermoFisher Scientific), whereas RNA integrity (RIN>7) was confirmed using the Agilent 2100 Bioanalyzer (Agilent, Santa Clara, CA).

### TISSUE DISSOCIATION FOR SINGLE-CELL ANALYSIS

Tissue was received fresh from the Brigham and Women’s Tissue and Blood Repository Core in cold RPMI-1640 medium (Gibco 11875093) with 10% fetal bovine serum, hereafter referred to as RP10). Research personnel were privy to preoperative biopsy histology, but otherwise masked to clinical information. All efforts were made to minimize time between tumor excision and single-cell processing. Tumor lysis was performed using the Miltenyi Tumor Dissociation Kit (Miltenyi Biotec, 130-095-929) per manufacturer guidelines. Tumors were sharply lysed on ice using Noyes spring scissors until fine enough to pass through a 2ml serological pipette. Tumors were then transferred to enzymatic media, mechanically dissociated using the gentleMACS tumor-02 protocol, and incubated for 15 minutes under continuous rotation at 37°C. Dissociation was repeated using the gentleMACS tumor-03 protocol and a second 15-minute incubation was performed. The tumor lysates were filtered through a 70uM filter and washed with cold RP10. RBC lysis was performed using ACK lysing buffer (Quality Biological, 118-156-101) per manufacturer protocol. Following an additional wash with 5ml PBS (Gibco), cells were resuspended in RP10, placed on ice, and assessed for concentration and viability using the Bio-Rad TC20 Automated Cell Counter per protocol. Epithelioid samples with mean viability below 70% were excluded; however, this viability threshold was not employed for biphasic or sarcomatoid samples to maximize collection of rare tumors. Samples of any histology were excluded if mean cellularity was less than 50,000 cells per mL.

### SINGLE-CELL RNA-SEQ (SCRNA-SEQ) DATA LIBRARY GENERATION, SEQUENCING, AND ALIGNMENT

ScRNA-seq processing followed the Seq-Well protocol, a method uniquely compatible with low-input samples (Gierahn et al., 2017). Briefly, on the day of processing, arrays were loaded to saturation with RNA capture beads (ChemGenes) and temporarily stored in bead loading buffer (100 mM sodium carbonate [Sigma 223530] with 10% bovine serum albumin [Sigma A9418]). Prior to cell loading, arrays were resuspended in 5 mL RP10. After dissociation, single-cell suspensions were counted as described above. Suspensions with adequate cellularity were diluted to a target range of 10,000-20,000 cells per 200 μL of RP10. To generate technical replicates in select samples, appropriately diluted single cell suspensions were divided into 200ul aliquots and loaded onto multiple separate arrays. These arrays then proceeded through the remainder of the Seq-Well protocol, including sequencing, in parallel. Excess RP10 was aspirated from the array and cells were loaded onto arrays. Excess cells were washed off with PBS (4×5 mL, Gibco), briefly left in RPMI (5 mL) and cell+bead pairs were sealed for 40 minutes at 37°C using a polycarbonate membrane (Fisher Scientific NC1421644). Arrays were rocked in lysis buffer for 20 minutes and RNA was hybridized onto the beads for 40 minutes. Beads were removed from the arrays using centrifugation and reverse transcription was performed overnight using Maxima H Minus Reverse Transcriptase (Thermo Fisher EP0753). Prior to sequencing, the beads underwent an exonuclease treatment (New England Biolabs M0293L) and second strand synthesis *en masse* followed by whole transcriptome amplification (WTA, Kapa Biosystems KK2602) in 1,500 bead reactions (50 μL). cDNA was isolated using Agencourt AMPure XP beads (Beckman Coulter, A63881) at 0.6X SPRI (solid-phase reversible immobilization) followed by a 1X SPRI and quantified using the Agilent 2100 Bioanalyzer system.

Library preparation was performed using Nextera XT DNA tagmentation (Illumina FC-131-1096) and N700 and N500 indices specific to a given sample. Tagmented and amplified sequences were purified with a 0.6X SPRI. cDNA was loaded onto either an Illumina Nextseq (75 Cycle NextSeq500/550v2 kit) or Novaseq (100 Cycle NovaSeq6000S kit, Broad Institute Genomics Platform) at 2.4 pM. The paired end read structure was either 20 bases (containing 12 bp cell barcode and 8bp UMI) by 50 bases (transcriptomic information) or 26 bases by 57 bases with an 8 base pair (bp) custom read one primer for NextSeq500 runs. For Novaseq runs, the paired end read structure for libraries was either 28 bases by 85 bases or 150 bases by 150 bases with either an 8bp or 10bp custom read one primer. The demultiplex and alignment protocol was followed as previously described (Macosko et al., 2015). While Novaseq data were directly output as FASTQs, Nextseq BCL files were converted to FASTQs using bcl2fastq2. The resultant Nextseq and Novaseq FASTQs were demultiplexed by sample based on Nextera N700 and N500 indices. Reads were then aligned to the hg38 transcriptome using the DropSeq tools (Macosko et al., 2015)pipeline implemented on the Brigham and Women’s ErisOne cluster using standard settings.

### BULK RNA-SEQUENCING ANALYSIS

Eighty-nine of the 93 samples had sufficient tissue to allocate for parallel bulk RNA-sequencing. Of these, 71 samples were adjacent to scRNA-seq sample. Another 11 and 7 were isolated from the same site or case, respectively. Extracted RNA was shipped to Medgenome on dry ice. Polyadenylated RNA were enriched using poly-T oligo attached magnetic beads. Reverse-stranded bulk cDNA libraries were prepared using the Illumina TruSeq stranded mRNA kit (#20020595) and sequenced to approximately 100 million 100bp paired-end reads per samples on the Novaseq 6000. Bulk RNA-sequencing reads examined for adequate mapping rates and library sizes. Quality-controlled libraries were then mapped to the human genome (hg38) and read counts were generated using STAR v.2.7.3 (Dobin et al., 2013). The resulting counts were normalized to transcripts per million (tpm) with a library size adjusted using trimmed moving means normalization in ‘edgeR’ (Chen et al., 2014; Robinson et al., 2009) for subsequent visualization and analysis. Additionally, the raw counts matrix was intersected with the raw counts matrix of 211 previously published PM transcriptomes and were assigned into the four reported consensus clusters using the 400 most variable genes identified in the previously published 211 transcriptomes as previously described (Bueno et al., 2016).

### HISTOPATHOLOGIC ANALYSIS

Eighty-two of the 93 samples had sufficient tissue to allocate an adjacent piece to histopathological sampling. For seven of the samples without adjacent tissue, slides were sectioned from tissue obtained from the same surgical resection; one of these was taken from the same anatomical site as the scRNA-seq sample. Four samples did not have sufficient tissue for histopathologic analysis. All slides were read by a licensed pathologist for relevant parameters including slide histology, composition (percent malignant cell, sarcomatoid, epithelioid, and fibroblast content), inflammatory score (0-3), nuclear pleomorphism score (0-3), notation of any non-pleural tissue observed, and whether sufficient tissue was present for analysis.

### BULK CNV IDENTIFICATION USING OPTICAL GENOME MAPPING

Banked fresh frozen tissue from each PM case was shipped on dry ice to Bionano for optical genome mapping assay and analysis. There, ultra-high-molecular-weight-DNA was extracted, followed by labeling, linearization, and imaging of DNA (Saphyr, Bionano). The results were analyzed using the Bionano Access Version 1.6 and 1.7 Rare Variant (>5000 bp) and Copy Number (>500,000 bp) pipelines and compared against the Bionano control database of 200 healthy individuals to exclude common germline CNVs. Resulting CNVs were examined for overlap with coding regions of PM associated genes. For each sample, base pair coordinates of observed CNVs were mapped to gene regions with sufficient scRNA-seq expression for inferred CNV analysis and compared to scRNA-seq CNV events.

### SNV IDENTIFICATION USING BULK WHOLE EXOME SEQUENCING

Whole exome sequencing (WES) was performed on paired bulk tumor and normal (blood) samples for each case. Exome capture was performed using Agilent SureSelect XT HS2 kit (#G9983A) targeting specific genomic regions (Human Exome V7 #5191-4005). Resulting libraries were sequenced to either 30X (normal DNA) or 100X (tumor DNA) depth with 100bp paired-end on the Novaseq 6000.

Reads from multiple lanes were uploaded to the Seven Bridges Cancer Genomic Cloud (www.cancergenomicscloud.org). Sequencing libraries from each case were concatenated using the Seven Bridges SBG FASTQ Merge CWL1.0 tool. Next, tumor-normal pairs were analyzed for somatic SNVs, copy number variants and tumor purity using the Seven Bridges “WES Tumor-Normal with Variant Calling, CNV estimation, TMB, MSI and HRD scores” public application (sgb:draft-2, v1.0) developed by PDXNet (https://cgc.sbgenomics.com/public/apps) using default parameters. Briefly, the WES tumor-normal pipeline utilized the Broad Institute’s best practices. Alignment was performed using BWA (Li & Durbin, 2009) to the hg38 genome. Variants were called with Mutect2 (Version 4.1.3.0) (Benjamin et al., 2019) and resulting variant calls were analyzed and annotated using the ‘SnpEFF’ (Cingolani, Platts, et al., 2012) and ‘SNPSift’ (Cingolani, Patel, et al., 2012) tools. The resulting somatic SNV calls are summarized with the imputed malignant cell and normal allele frequencies, the gene impacted, and the mutation effect rank (High, Moderate, Modifier, or Low) in **Table S1C**. CNVs were also identified, and tumor purity and loss of heterozygosity (LOH) regions were estimated using the ‘Sequenza’ R package (Favero et al., 2015) by assessing the variation of tumor/normal read coverage and minor allele frequencies across the exome-targeted genome.

To minimize false positives due to low allele frequency sequencing errors and numerous C>A/G>T calls, the somatic mutations detected with “high confidence” were required to have an imputed tumor allele frequency > 0.1 and tumor read coverage of at least 10 reads. Somatic mutations in the most mutated genes in PM were extracted (**Figure 1B**; **Table S1C**).

### SINGLE-CELL DATA QUALITY PRE-PROCESSING AND INITIAL CELL TYPE DISCOVERY

All single-cell data analysis was performed using the R language for Statistical Computing (v4.0.5). Digital gene expression (DGE) matrices (cells x genes) containing UMI expression values of genes in 10,000 “cells” for each sample or replicate were initially aggregated by batch to assess distribution of genes, detected, UMIs, and mitochondrial reads for bimodality (suggesting natural threshold for information poor and rich cell observations). After inspection and threshold determination, each sample or replicate’s DGE was then filtered to exclude low-quality cells (<80 genes detected; <300 UMIs; >40% mitochondrial reads) before being merged (preserving all unique genes) to create a DGE containing all 114 samples and replicates. The merged dataset was further trimmed to remove cells with >10,000 genes which represented outliers and likely doublet cells. Genes that were not detected in at least 3 cells were also removed. The resulting DGE matrix contained 279,154 cells.

Using the R package Seurat (v4.0.1) (Satija et al., 2015), PCA was performed over 3,000 variable genes normalized using SCTransform (Hafemeister & Satija, 2019). Additionally, UMI count data for 15,000 genes were normalized SCTransform controlling for mitochondrial content (Hafemeister & Satija, 2019) to yield normalized counts and log-counts for downstream data visualization and exploration of additional genes not utilized in clustering. After examining the distribution of variance across components, 110 PCs were input to build an SNN graph and cluster cells (res=2; k.param=20). This identified clusters shown using UMAP (n.neighbors=30) in **Figure S1A**. Lower quality cells (e.g. mitochondrial reads >25% but <40%) were included under the expectation that cells primarily defined poor quality would cluster together (**Figure S1B**). Thus, 6,204 cells from cluster 17 were removed because they only expressed mitochondrial markers and 589 cells from cluster 70 were also removed due to their low UMI to genes expressed ratio (likely multiplets). Doublet cells were then identified using scDblFinder (Germain et al., 2021) and subsequently removed (n=6,678). Notably, clusters 6 (n=9,214) and 61 (n=1,046) were flagged but retained as they simultaneously expressed high mitochondrial UMIs and clear cell type markers (**Table S2**). This led to an overall high-quality dataset of 266,265 single cells assigned to 83 clusters with low overall fraction of mitochondrial reads (median =0.091) for downstream analysis (**Figure S1C**).

The expression of known markers was used to collapse clusters containing shared lineage information in the trimmed dataset. For example, clusters 1, 15, 28, 32, 34, and 81 all express high levels of CD8+ T cell markers—*CD3D*, *TRBC2*, and *CD8A*—and were accordingly collapsed for this first pass analysis (**Figure S2A**). To aid cell type identification, the Wilcox test implemented in Seurat was performed to identify top marker genes in each cluster (**Table S2**).

### SINGLE-CELL CNV IDENTIFICATION

Single-cell CNVs were estimated as previously described by computing the average expression in a sliding window of 100 genes within each chromosome after sorting the detected genes by their chromosomal coordinates (Patel et al., 2014; Tirosh, Izar, et al., 2016; Tirosh, Venteicher, et al., 2016). While all 266,262 cells were readily assigned to phenotypic groups, CNV identification was performed on a subset (136,910 cells) consisting of the highest quality cells (at least 400 genes expressed and 750 UMIs detected). Thirty cells were of sufficiently high quality but could not be analyzed using inferCNV due to limited phenotypic representation (cluster size = 1) in their respective surgical case. For this analysis, all NK, CD45+ proliferating, Epithelial, EM hybrid, Mesenchymal, Mural, and Endothelial cells identified as above, of sufficient quality, and isolated from three normal pleura samples were utilized as reference normal populations (n=422). Complete information on the inferCNV workflow used for this analysis can be found at the Broad Institute GitHub (https://github.com/broadinstitute/inferCNV/wiki).

Initially, CNVs were detected in all 136,880 cells of sufficient quality. As previously described (Tirosh, Izar, et al., 2016), specialized immune cells such as B cells, monocyte, and macrophages exhibited heterogeneous CNV profiles due to their chromatin state, so these cells along with CD4+ T, CD8+ T, and Treg cells (n= 72,322 cells) were annotated as immune cells and excluded from further CNV analysis. For reference and candidate malignant cells (n=64,558), these CNVs were utilized to compute CNV score and correlation of CNV profile with the average CNV signal for the top 5% of altered cells in each surgical case as previously described (Tirosh, Izar, et al., 2016). Next, a subset of cells was selected with the purpose of capturing putative malignant cells to enable computationally feasible subclone analysis. Cells were first separated into non-malignant and putative malignant by entropy of patient distribution for their respective phenotypic cluster (see above; **Figure S3A-B**). Visually determined thresholds using CNV score as a function of CNV correlation divided cells into “tumor”, “ambiguous”, and “not tumor” categories for each surgical case (**Figure S3C**). Cells annotated as “tumor” or “ambiguous” were considered suspicious for malignancy and included in subclonal analysis. Additionally, all cells within clusters where greater than or equal to 60% cells, including those annotated as “not tumor”, were considered suspicious for malignancy were also included in subclonal analysis.

### SUBCLONAL ANALYSIS WITH SINGLE-CELL INFERRED CNVS

For cells considered suspicious for malignancy and of sufficient quality, subclonal copy number variants were called with high sensitivity using the inferCNV workflow comprehensively outlined at the Broad Institute GitHub (https://github.com/broadinstitute/inferCNV/wiki) (Fan et al., 2018; Patel et al., 2014; Tirosh, Venteicher, et al., 2016). Identical reference cells from control cases were utilized as in the initial CNV identification. Briefly, a six-state Hidden Markov Model (i6-HMM) was used to predict relative copy number status (complete loss to >3x gain) across putative altered regions in each cell. A Bayesian latent mixture model then evaluated the posterior probability that a given copy number alteration is a true positive. The default cutoff (BayesMaxPNormal = 0.5) was used for this step. The results of this filtered i6-HMM output were then used to cluster the single cells using Ward’s method. Statistical significance of potential subclusters was tested using the “random trees” method (*P* < 0.05, 100 random permutations for each split) implemented in inferCNV. Only significant subclusters were retained. To track subclonal heterogeneity across anatomic sites, the above workflow was implemented for each surgical case including all samples and replicates allowing the CNVs to determine cell sorting agnostic to sample-of-origin (**Figure S4**).

In clonal CNV profile comparison analysis, inferred CNVs of each clone were mapped to chromosomal bands. Band-summarized CNV profiles were then hierarchically clustered using the Manhattan distance and ward.D2 linkage method (**Figure 1E**).

### IDENTIFICATION OF MALIGNANT CELLS

Malignant cells were identified through an iterative process combining cluster phenotype, patient-cluster distribution (entropy group), detected CNVs, and clonotype. After excluding clearly non-malignant immune cells (e.g., macrophages, T cells, etc.), candidate malignant cells were identified by examining the distribution of cells within each phenotypic cluster across patients operationalized by normalized Shannon entropy (**Figure S2B**); cells in clusters predominantly identified in a single patient (low entropy) were considered ‘putative tumor’ (**Figure S2C & S2G**). Next, the CNV signal score and correlation to the average profile of the top 5% altered cells were computed for each surgical case and utilized to enrich for cells suspicious for malignancy as described (see ‘Single Cell CNV Identification’) with the intention of using subclonal analysis to identify remaining non-malignant cells.

For 52,937 (82.0%) of the 64,558 tested cells, entropy group and CNV metric malignant cell annotations were concordant (**Figure S2D; Table S2**). The 1,200 cells that were not considered suspicious for malignancy, but putatively malignant by Shannon entropy analysis were either isolated from non-malignant control samples (72 cells) or found within cluster 35 (1,128 cells). The profiles for the discordant cluster 35 cells were examined in the 6 surgical cases containing the most cluster 35 cells (**Figure S4**) revealing these cells to be non-malignant. A total of 15,212 cells (9,054 stroma and 6,158 immune) with CNV analysis were concordantly defined as “non-tumor” by entropy group and CNV metrics. Subclonal analysis was performed on the remaining 49,346 cells suspicious for malignancy. Assigned clones were examined for evidence of CNVs for each patient and assigned to “malignant” or “non-tumor” accordingly (**Figure S5**). Some clones were partitioned on entropy group, detected CNVs, and phenotype due to visually obvious intraclonal discrepancies in CNV evidence. In these cases, individual partitions were assigned to “malignant” or “non-tumor”. Thus, 64,558 cells were characterized as “malignant” (40,501 cells) or “non-tumor” (16,834 stroma and 7223 immune). Among “malignant” cells, 1,303 (3.22%) clustered and expressed markers consistent with stromal or immune identity (**Figure S2E)** and another 3,702 expressed *PTPRC*/CD45; these were excluded as likely doublets leaving 35,496 malignant cells (**Figure S2F & S2G**).

Strictly for the purposes of tumor micro-environment description and cellularity computation, malignancy was inferred from phenotype and entropy group alone for the remaining 129,352 cells with insufficient data for CNV analysis. Notably, entropy group alone was sensitive (90.90%) and specific (86.93%) for malignant cells in the data where CNV analysis was available.

### SUBCLUSTERING OF MALIGNANT AND NON-MALIGNANT CELLS

Detailed phenotyping required splitting the dataset into malignant and stroma (non-immune, non-malignant) fractions and restricting analysis to the highest information cells (greater than 400 genes and 750 UMIs). The 945 PNS malignant cells were excluded leaving 34,551 PM malignant cells. After sub-setting to only malignant cells, initial PCA was performed using the first 35 PCs on the re-scaled data for SNN clustering (resolution = 1.5). Several low-variance dimensions of this initial PCA had significant contributions from mitochondrial gene features and 2 malignant cell clusters contained cells with high mitochondrial reads (first quartile >25%). Cells in these clusters were removed in subsequent analysis leaving 32,383 high quality PM malignant cells. As with malignant cells, PNS stroma (859 cells) and CD45+ cells (3,214) leaving 12,776 high quality PM stromal cells (no high mitochondrial stromal clusters identified on initial PCA). PCA was repeated on filtered malignant or stromal cells using the first 25 PCs for SNN clustering (resolution=1.2) and UMAP visualization.

Marker genes were used to identify previously described stromal cell types (**Figures S10; Table S5A**). Because the previous malignant cell identification emphasized specificity not sensitivity, stroma clusters were assessed for patient-specificity to maximize enrichment of non-malignant cells for further analysis. Clusters where greater than 90% of cells arose from a single patient were considered patient-specific clusters and excluded from downstream stroma phenotyping. These were stroma clusters 33 (100% from 27409), 8 (96.9% from 27306), 23 (96.8% from 27444), 0 (96.3% from 27409), 21 (93.0% from 27444), 32 (91.9% from 27042), and 1 (91.9% from 30285). Clusters 6, 16, 17, 19, 22, 23 expressed lineage markers from multiple immune and stroma populations; these were likely doublets or cells with ambient RNA in the setting of necrosis. A total of 4,566 cells were excluded from downstream variation and malignant vs stromal cell differential expression analyses, where 2,695 cells were found in patient-specific clusters, 2,093 cells from clusters expressing multiple lineage markers, and 222 cells in cluster 23 with both features.

For immune micro-environment comparisons and lineage signatures, analysis was also restricted to 79,552 immune cells with greater than 750 UMIs and 400 expressed genes. Furthermore, immune cells of lymphoid (“CD8+ T cell”, “CD4+ T cell”, “Treg”, “B cell”, “CD45+ proliferating”) and myeloid (“Macrophage”, “Monocyte”, “NKC”, “Mast cell”) origin were filtered to contain less than 10% and 15% mitochondrial reads, respectively, leaving 64,163 immune cells for signature and TME association analysis. PCA was repeated on filtered immune cells using the first 35 PCs for SNN clustering (resolution 1.2) and UMAP visualization. Clusters were assigned into broad immune types (**Figure S11; Table S6**). Several clusters contained at least 20% cells expressing non-immune sarcomatoid and epithelioid genes, e.g. *ITLN1, COL3A1*A. These clusters were annotated as “Multiplets/Ambient” and excluded from further analysis. Additional sub-clustering analysis of NK/T-cells (30 PCs, resolution 1.2) and macrophages/dendrocytes (30 PCs, resolution 1.2) were performed facilitating more granular functional sub-type assignment (**Figure S11D-E**). For macrophages/dendrocytes, as previously observed (Raghavan et al., 2021; Zhang et al., 2020; Zilionis et al., 2019), initial clustering identified a cluster of macrophage-T-cell doublets. Additionally, the first principal component was driven by heat stress response expression and three clusters contained at least 75% of cells expressing heat stress response genes above the median. These four clusters were labelled appropriately, and the remaining 16,922 cells were re-clustered as above. Clusters predominantly of macrophage-T-cell doublets and heat shock response expression were also identified and removed in NK/T-cells, resulting in 24,206 cells clustered as previously described.

Subsequent phenotyping for malignant, stromal, and immune cells is discussed below (**Generation of expression signatures/scores**).

### GENERATION OF SINGLE CELL EXPRESSION SIGNATURES & SCORES

The CV score (Bueno et al., 2016; Severson et al., 2020) for each cell was computed by dividing scTransform normalized UMI counts of VIM+0.01 from CLDN15+0.01 and performing a base two logarithm. All other expression scores were computed as previously described by taking a given input set of genes and comparing their average relative expression to that of a control set (n=100 genes) randomly sampled to mirror the expression distribution of the genes used for the input (Raghavan et al., 2021; Tirosh, Izar, et al., 2016). While all scores were computed in the same way, choosing the genes for input varied. The relevant approaches have been outlined below. Where correlations (Spearman’s *r)* are performed over genes, we used the log-transformed scTransform normalized UMI count data for each case.

#### Previously Published PM Bulk RNA-Seq Signatures

All gene sets were filtered for genes detected (normalized UMI >=1) in at least three malignant or stroma cells. Parenthetical E and S were added to gene set names to indicate similarity to Blum E- and S-score where appropriate. E-score and S-score gene sets consisted of the detected genes found within the E-score and S-score components as reported by Blum *et al* (Blum et al., 2019). The four continuous scores reported by Alcala *et al*, i.e. angiogenesis-low (E), angiogenesis-high (S), inflammation-high, inflammation-low, were generated by taking the 30 detected genes with the highest positive or negative correlation with the relevant PC as previously published (Alcala et al., 2019). The two TCGA iCluster signatures were generated by taking the top 30 genes ordered by cluster centroid scores of the 2,807 mRNA classifier genes for mRNA iCluster 1 (E) and mRNA iCluster 3 (S), respectively (Hmeljak et al., 2018). The microarray derived C1 (E) and C2 (S) signatures were generated by identifying the subset of the published 40 discriminating genes (Reyniès et al., 2014) that were detected in the single cells and significantly different (t-test, p-value<0.05) between C1 and C2 and then assigning genes with higher expression to the appropriate cluster signature. Gene sets are reported in **Table S2A**.

#### Cell Cycle

The previously established signatures for G1/S (n=42 of 43 genes) and G2/M (n=55 of 55 genes) were filtered for genes detected in malignant cells and utilized to assess cycling status (Tirosh, Izar, et al., 2016). After inspecting the distribution of scores in the complete dataset, any cell >1.5 s.d. above the mean for either the G1/S or the G2/M scores were annotated as cycling (Galen et al., 2019).

#### mSigDB Signatures

Relevant gene sets were retrieved from mSigDB and GSEA analysis was performed as below. Scores were computed using the union of unique leading-edge genes identified in each respective gene set (**Table S3C**).

#### scRNA-seq Malignant Cell Derived E-scores and S-scores

All detected genes were correlated (Spearman) with the relevant histomolecular-related PC 1. Candidate genes (|*r*| > 0.025) observations were scrambled in 1000 iterations to generate an empiric expression-normalized gene-specific null of correlation to the PC. Genes with empiric q-value ≤ 0.0001 and *r* ≥ 0.15 or r ≤ −0.15 formed the scRNA-seq S-score (tS-score: 232 genes) and E-score (tE-score: 371 genes) signatures, respectively (**Tables S3E**). Scores were computed and visualized using the top 30 genes in each category (**Figures 2B & S7A**).

#### ECM-A, ECM-B, ECM-A.27306, and ECM-B.27306

To minimize EM gradient signal, differential expression was performed comparing the top and bottom deciles of PC 2 (ECM variation) within the bottom (Epithelioid-like) and top (Sarcomatoid-like) quartiles of PC 1 (**Table S4A**). For ECM-A and ECM-B signatures, the resulting genes with adjusted p-value < 0.01 and average log2 fold-change ≥ 0.25 or ≤ −0.25, respectively, from each PC1 quartile were intersected to identify a list of ECM genes unrelated to EM gradient. The genes meeting identical criteria in the top quartile (Sarcomatoid-like) of PC 1 formed the ECM-A.27306 and ECM-B.27306, which were created to describe the ECM variation primarily captured by the case 27306. Resulting gene signatures are reported in **Table S4C.** All signature genes were utilized for computing score, but only top 30 were used for visualization (**Figure S7E**).

#### Histomolecular Commitment Distance and Score

A subpopulation of malignant cells equivalently expressing low epithelioid and sarcomatoid programs was revealed when cells were ordered by EM gradient (**Figure S8A-B**). To characterize these malignant cells without histomolecular commitment (HC), the HC distance’ was computed by computing the Euclidean distance from equal expression of tE-score and tS-score signatures for each tumor cell (**Figure S8C**). Cells with an HC distance 1.5 standard deviations below the mean HC distance, i.e. less than 0.94, were considered uncommitted. To generate an HC score, all detected genes were then correlated (Spearman) with computed Euclidean distance to generate a HC correlation. To enrich for genes specific to uncommitted cells, the absolute value of each genes’ correlation to PC 1 was added to their HC correlation resulting in a corrected HC correlation. In this way, genes highly correlated with tS-score or tE-score signatures were excluded from the uncommitted (U) score. Several genes with low corrected commitment correlation were ubiquitously expressed; to filter these genes, differential expression between uncommitted cells and epithelioid or sarcomatoid committed populations was performed (**Table 3F**). Genes with corrected commitment correlation ≤ −0.05 (empiric q-value ≤ 0.0001), significantly differentially expressed in both comparisons (adj. p-value <0.01), and expressed in less than 20% of cells in either committed population were considered uncommitted genes and formed the HC score signature (n=48 genes) (**Figure S8D; Tables S3E-F**). Cells were visualized and scored using the 30 genes with the greatest negative corrected commitment correlation (**Figures S8C & 3D-E**).

#### Inflammation Score

All detected genes were correlated (Spearman) with the relevant inflammation-related PC 3. Genes (n= with empiric q-value ≤ 0.0001 and *r* ≥ 0.15 formed the scRNA-seq inflammation score (Inf-score) (**Table S3E,G**). All signature genes were utilized for computing score, but only top 30 were used for visualization (**Figure 2C**).

### PATHWAY ANALYSIS WITH GSEA AND SELECTION-UNBIASED CATEGORY ENRICHMENT

For gene set enrichment analysis (GSEA), the gene set search space consisted of canonical pathways and hallmark gene sets from MSigDB (Liberzon et al., 2011, 2015) and previously published PM gene sets as described above. These were utilized for GSEA using ‘fgsea’ (Korotkevich et al., 2019), where the enrichment metric was computed from either ordered embeddings of each PC dimension or average log2 fold-change in gene expression. For GSEA on PC embeddings, general biological perturbations for each component were inferred where possible (**Figure S6B**) from resulting significant gene sets (adj. p-value <0.05) reported in **Table S2B**. Significant gene sets (adj. p-value<0.05) resulting from GSEA of log2 fold-changes in gene expression between top and bottom deciles of PC 2 are reported in **Table S4B**.

For selection-unbiased category enrichment, over-representation of KEGG pathways was tested accounting for gene length for relevant gene signatures using ‘goseq’ (Young et al., 2010). Genes without KEGG pathway categories were excluded and the over-representation scores were computed with the “Wallenius” method. Pathways were considered significant if FDR ≤ 0.5. Pathway over-representation results are reported in **Tables S3B and S4D**.

### ASSIGNMENT OF PSEUDO-BULK MALIGNANT CELL STATE

Cases (n=29) and samples (n=64) with at least 10 malignant cells were assigned to “Sarcomatoid”, “Epithelioid”, and “Uncommitted” by averaging expression of the top 30 genes of each gene program across all malignant cells isolated from the case or sample. This generated a pseudo-bulk pure malignant cell case or sample with 90 gene features representing the three malignant programs (**Figures 3H & S12B**). The Pearson correlation distance was computed and used to hierarchically cluster with “Ward.D2” linkage cases or samples into three pseudo-bulk clusters representing the average malignant cell state found in the tumor tissue (Figures 3G & S12A). Individual cell-type composition of pseudo-bulk cases or samples was visually confirmed and corresponded to pseudo-bulk assignment (**Figures 3I & S12C**).

### TME ASSOCIATIONS

Samples were filtered for sufficient isolation of non-malignant cells (n > 200). Thus, of 89 samples, 29 were excluded for insufficient capture of TME cell types leaving 60 samples from 29 cases for TME association analysis. Of these, 15 samples had insufficient malignant cells (n < 10) to be assigned pseudobulk malignant cell type. The TME association with malignant cell phenotype was analyzed in two steps as previously described in pancreatic samples (Raghavan et al., 2021). The Simpson’s Index (a measure of ecological diversity) was computed using the count of each non-malignant cell type captured from each sample as input and correlated with each PM sample’s scRNA-seq malignant cell histologic score, i.e. tS-tE score (**Figure 4E**). In these samples, the number of non-malignant cells isolated was weakly, but significantly correlated with EM scRNA-seq malignant cell state (R^2^=0.087; p=0.0418) but not sample Simpson index (R^2^=0.053; p=0.0791). Next, the fractional representation for every non-malignant cell type in each PM resection sample was used to compute the pairwise correlation distance (Pearson’s *r*) followed by hierarchical clustering using Ward’s D2 method identifying TME relationships between samples. Because identified cell type assignment is dependent on choice in cluster granularity, stromal and immune cell types were clustered using Pearson correlation and Ward’s D2 clustering to identify relationships between identified cell types (**Figure S15A-B**). Features in this analysis were restricted to stroma marker genes (**Table S5A)** or immune, myeloid, and lymphoid marker genes (**Tables S6A,6C-D**), respectively.

Additionally, trends in fraction TME cell content by pseudobulk type were visualized by sample (**Figure S14C-D**). To identify significant associations between predominant malignant cell pseudobulk phenotype and TME components, distributions of fraction cells were compared for each cell type using a Kruskal-Wallis rank-sum test (**Figure S15C**). Values were considered significant for q-value <0.1, after correction for multiple hypothesis testing.

### GENERATION OF BULK RNA-SEQ SCORES FROM SINGLE CELL EXPRESSION

Bulk RNA-seq profiles obtained in this study as well as previously published (Bueno et al., 2016) were scored using single cell expression signatures above using single sample gene set enrichment analysis (ssGSEA) (Barbie et al., 2009; Subramanian et al., 2005). These scores were computed using the GSVA package in Bioconductor (Hänzelmann et al., 2013). Additional signature scores were generated from single-cell expression data to deconvolute bulk RNA-seq profiles, but not utilized to score individual cells as above. The generation of these signatures are described in detail below.

#### Stromal Subpopulation Signatures

Stromal subpopulation signatures were established using iterative differential expression (DEG) analysis analogous to the approach described by Krishna *et al* (Krishna et al., 2021). Initial, “subpopulation” DEG analysis compared each cluster (e.g. *TGFBI*+ fibroblast.1 cells) to all other cells in its broad lineage (e.g. all fibroblasts). This round of DEG establishes the DEGs specific to the cluster (i.e. subpopulation). For each cluster, only those genes (log FC > 0; adj. P < 0.01) exclusive to that cluster’s subpopulation DEG were retained. Subsequently, for each cluster a “lineage” DEG analysis, in which each cluster (i.e. *TGFBI*+ fibroblast.1 cells) was compared to all other clusters of other lineages (e.g. all mural cells, mesothelium, endothelium, *etc.*). For each subpopulation (e.g. *TGFBI*+ fibroblast.1, mesothelial cell.1, etc) within a lineage (e.g. Fibroblast, Mesothelium, Mural, Alveolocyte, Endothelium, Erythrocyte), the same set of lineage genes, i.e. the set of genes that are commonly upregulated across at least 70% subpopulations of the same lineage when performing DEG analyses vs other lineages were retained. To optimize lineage specificity, only genes that were expressed in less than 20% of subpopulations of other lineages were retained in signature. To define the final signature genes for each cluster, its subpopulation and lineage DEGs were combined (**Table S5C**). In parallel, all genes were identified that were present in at least 20% of subpopulations within a lineage in either subpopulation or lineage analysis. These genes represent marker genes for stroma lineages and subpopulations which may distort bulk estimates of malignant cells (see *Malignant-exclusive E- and S-scores).* To ensure that genes in each signature were only expressed in non-malignant cells, malignant cells were also included in analysis. Each malignant cell was assigned a lineage of either Sarcomatoid, Uncommitted or Epithelioid and a subpopulation category defined by its lineage and patient of origin (e.g. ‘Sarcomatoid.27306’). In this way, malignant lineages and subpopulations provided filters for stromal signatures.

#### Immune Subpopulation Signatures

To identify immune infiltrate contributions to previously described bulk PM gene signatures, PM immune infiltrate signatures were established for subpopulation and broad immune phenotype (e.g. Mast cell, Treg, NKC, B cell, Monocyte, CD4+ T cell, CD8+ T cell, and Macrophage). Immune subpopulations were defined by initial phenotype clustering (**Figure S2A**) and signatures were defined using an analogous approach to the generation of stromal subpopulation signatures. Treg clusters were considered part of the CD4+ T cell lineage for this analysis.

#### Malignant-exclusive Uncommited, E- and S-scores

To deconvolute bulk samples, scRNA-seq tE-score and tS-score signatures derived from malignant cells were filtered to exclude previously detected stroma marker genes (**Table S5D**). Additionally, the stroma marker gene-filtered tS-score signature was filtered to only include genes significantly differentially expressed that were also expressed in twice as many sarcomatoid cells when compared to fibroblasts (**Table S5F**) to generate the malignant cell exclusive sarcomatoid signature (meS-score: 13 genes). Given the absence of distinguishing genes separating mesothelium and epithelioid, no additional filtering following stroma marker gene exclusion was performed to generate the malignant cell exclusive epithelioid signature (meE-score: 91 genes). For the malignant exclusive uncommitted score (meHCscore), no differential expression filtering was applied resulting a 24 gene signatures (**Figure S14A-B; Table S5D-E**).

### IDENTIFICATION OF CANDIDATE TME EM GRADIENT DRIVERS

To identify factors in the TME that drive EM cell state in either the malignant cells themselves or in associated non-malignant cells, differential expression analysis was performed on a short-list of candidate EM driving ligands. Candidate ligands were identified in MSigDB (Liberzon et al., 2011) annotated with the “signaling receptor binding” molecular function gene ontology excluding immunoglobulin complex genes (1,481 genes). A final list of candidate EM driving ligands (630 genes) was created by intersecting the signaling genes with 1,891 genes coding for secreted protein obtained from The Human Protein Atlas (Uhlén et al., 2015, 2019) by searching “protein_class:Predicted secreted proteins”. For autocrine factors, differential expression analysis was performed using these candidate ligands comparing malignant cells with the “Sarcomatoid” cell state to those with the “Epithelioid”, and then comparing “Uncommitted” malignant cells to the rest. A gene was considered differentially expressed if q < 0.05 and log(fold change) > 0.2 in either comparison. Genes were then assigned to subtypes based on the log fold change direction (**Figure 5D, Table S7A**). Paracrine factors were determined in a similar manner with slight modifications. Non-malignant cells were grouped into “Sarcomatoid”, “Epithelioid”, or “Uncommitted” based on clustering on average malignant cell expression of malignant programs in their respective tumor samples (**Figure S12**). Differential expression was then assessed between all cells from a given group contrasted to the remaining cells as above (**Figure S18A, Table S7B**). Differentially expressed genes were filtered by significant interactions identified using CellPhoneDB (**Table S7C**). This resulted in a short list of candidate factors expressed by either non-malignant (**Table S7D**) or malignant cells (**Table S7E**) in the PM microenvironment. Average expression for each factor was visualized in each cell type isolated from each of the respective malignant cell state-specific TMEs (**Figures S18B-C**).

