## Supplemental Figures for "Comprehensive multi-site profiling of the malignant pleural mesothelioma micro-environment identifies candidate molecular determinants of histopathologic type"

### Supplementary Figures

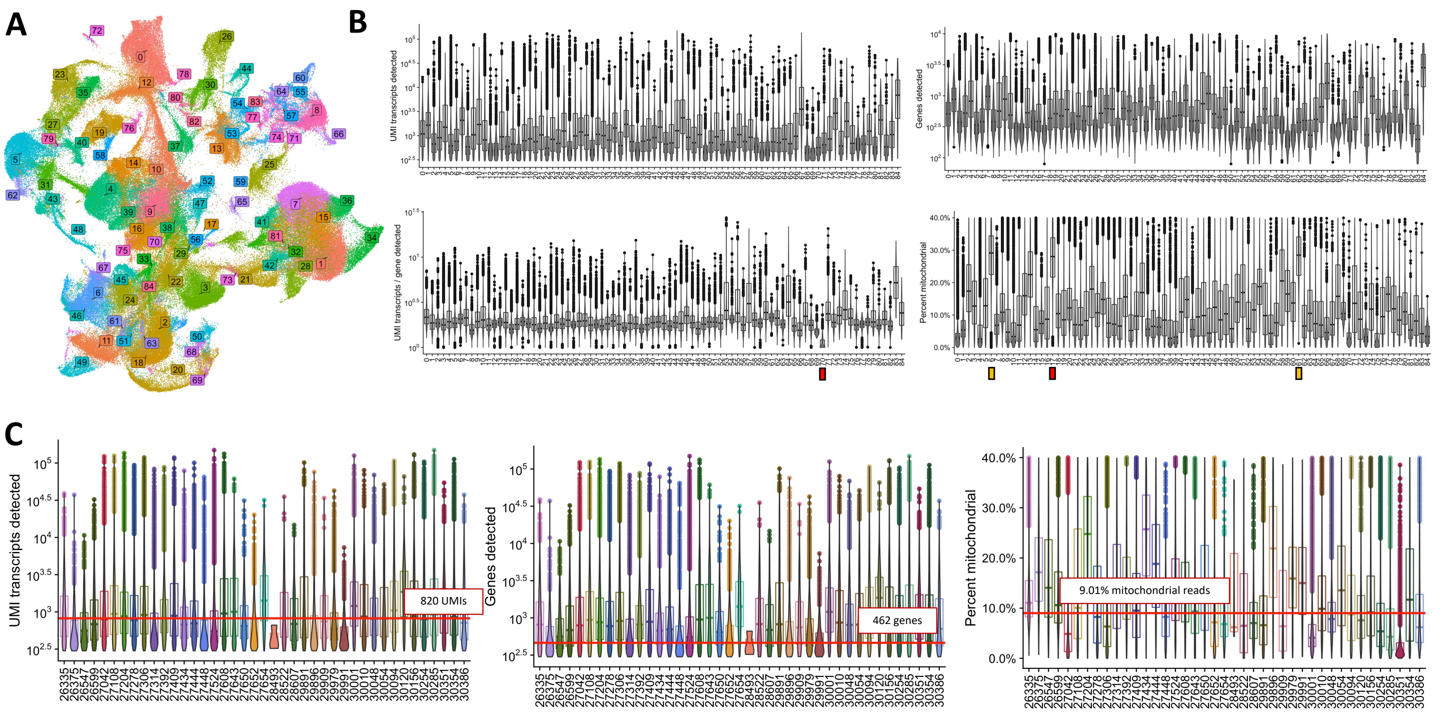


**Figure S1: Clustering single cells and quality filtering, related to Figure 1. A)** UMAP displays 279,154 cells colored by SNN clusters computed in Seurat. **B)** Boxplots with superimposed violin plots display distribution of UMIs detected (upper left), genes detected (upper right), ratio of UMIs to genes detected (lower left), and mitochondrial fraction (lower right), within each cluster. Clusters removed in quality control for a given parameter are denoted with a red box. Two clusters with high mitochondrial content, but easily identified using marker genes are denoted with a yellow box. **C)** Boxplots with superimposed violin plots display distribution of mitochondrial fraction, UMIs detected, and genes detected across patients (x-axis) for cells following additional quality filtering of doublets, mitochondrial content, and expressed UMIs/genes. Median values are indicated with text and horizontal intercept.

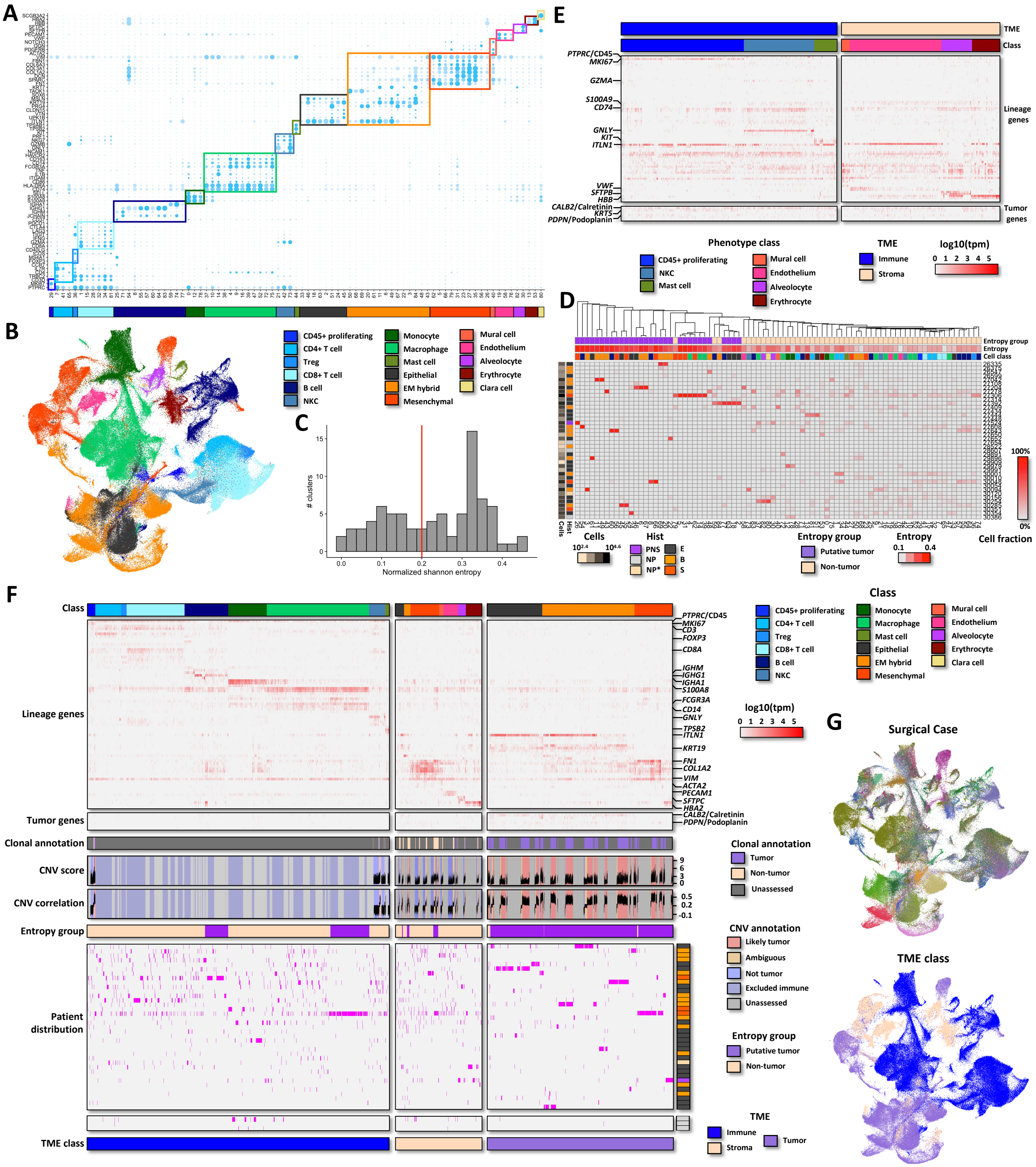

**Figure S2: Cell type and malignant cell identification, related to Figure 1. A)** Dotplot displays average expression (color) and fraction of cells expressing (size) of lineage genes across 83 clusters. Colored boxes denote gene programs associated with specific cell phenotypes assigned to each cluster annotated below. **B)** UMAP plot displays 266,252 cells projected into two dimensions and colored by cell phenotype class. **C)** Histogram displays the normalized Shannon entropy of cell distribution across patients in the 83 clusters. Vertical red line indicates threshold separating putative tumor from non-tumor clusters. **D)** Heatmap displays the fraction of cells isolated from each surgical case (rows) for each cluster (columns). Surgical cases are annotated with the number of cells isolated and diagnosis histology. Clusters are annotated with assigned entropy group, normalized Shannon entropy, and phenotypic class. **E)** Heatmap displays log10(tpm) expression of lineage and tumor genes of 1,303 immune and stroma phenotype cells assigned tumor by clonal analysis. Cells are annotated with tumor microenvironment and phenotype class. **F)** Multi-modal identification of tumor cells includes phenotype assignment using expressed genes (top), inferred CNVs (middle), and distribution of identified clusters across patients (bottom). **G)** UMAP plots display 266,252 cells projected into two dimensions and colored by case (top) or assignment into malignant, immune, and stroma cells (bottom).

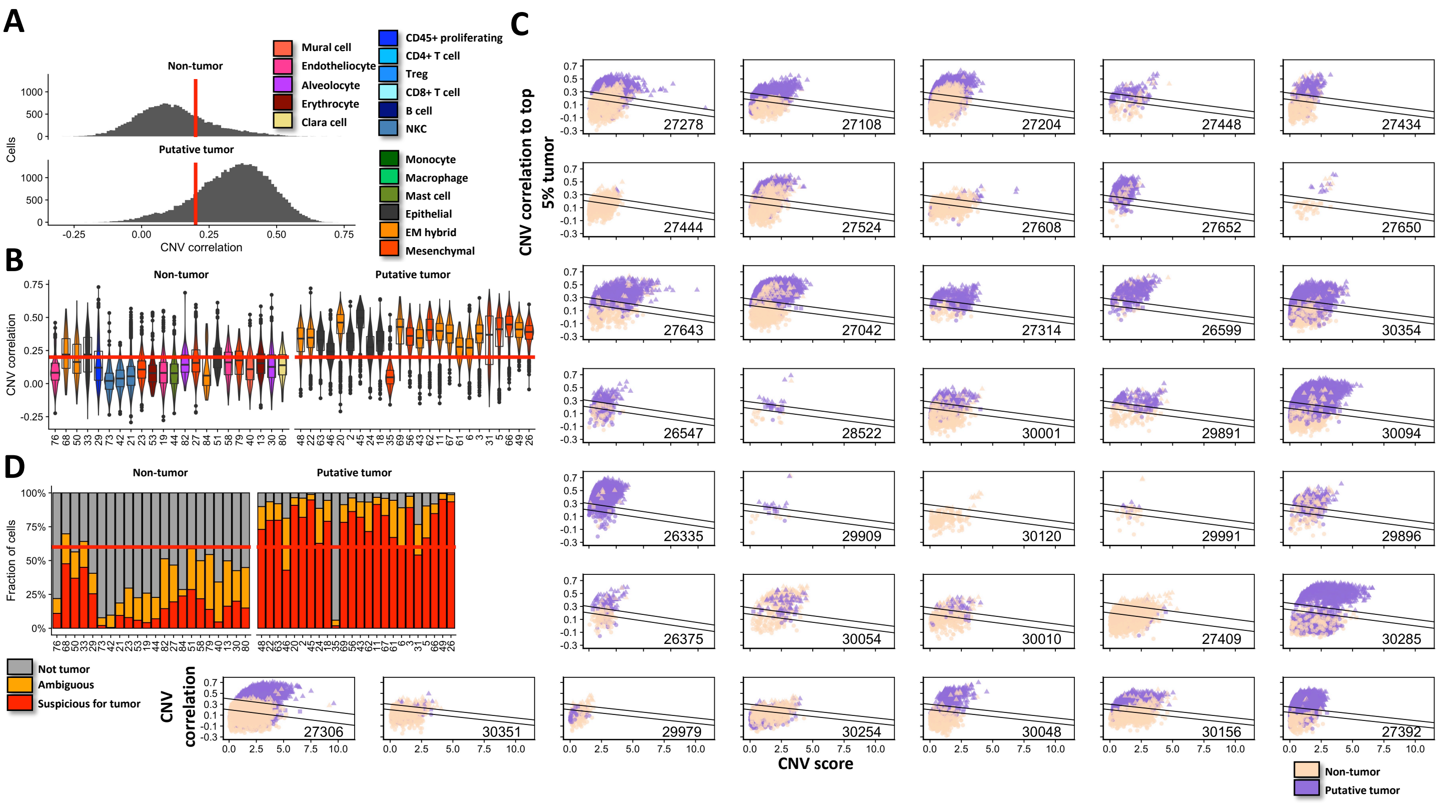


**Figure S3: CNV score and correlation to top tumor facilitate identify candidate tumor for clonal analysis, related to Figure 1. A-B)** Distributions of CNV correlation to top 5% tumor for non-tumor and putative tumor as determined by phenotype distribution across patients across whole population (histogram, **A)** or within individual phenotype clusters (Violin and boxplots, **B)**. Clusters are colored by molecular phenotype and ordered by normalized Shannon entropy. **C)** Each panel displays a scatterplot for a MPM or control case. Scatterplots display CNV score (x-axis) vs correlation to top 5% of tumor (y-axis). Candidate tumor status defined by entropy of phenotype cluster is indicated by color and designation as ambiguous, tumor, or non-tumor is indicated by shape. Lines in each scatterplot define thresholds utilized for tumor enrichment designation. **D)** Barplots display the distribution of assignment for the purposes of enriching tumor across phenotype clusters.

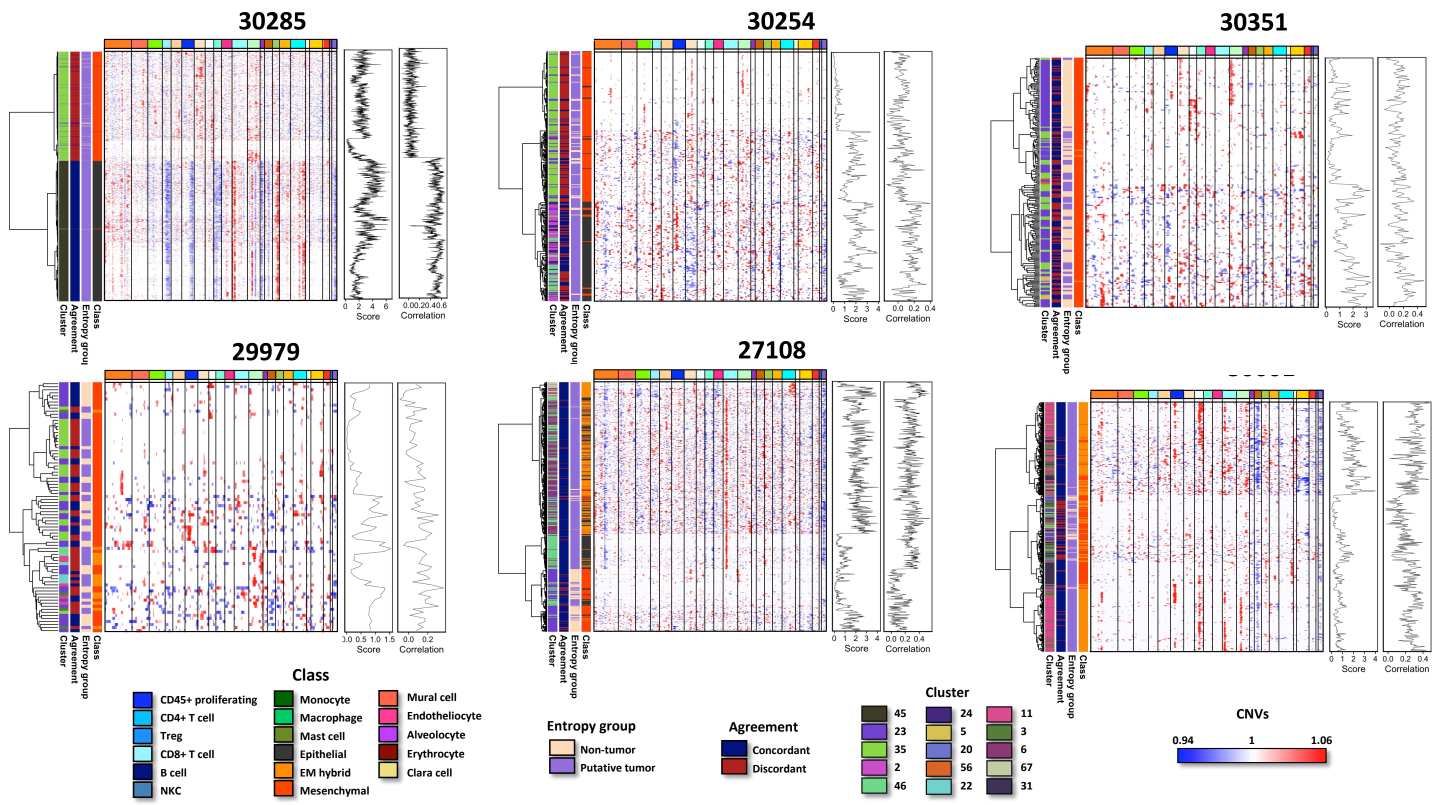


**Figure S4: Hierarchical clustering of inferred CNVs of select populations confirm that cluster 35 is non-malignant.** Panels display heatmaps hierarchically clustered CNV profiles of cells (y-axis) from select cell populations (cluster 35, with confident non-tumor, i.e. cluster 23, and case-specific tumor)**.** Cells are annotated with phenotype, cluster-distribution tumor classification, agreement between tumor classifications, and cluster. Line plots to the left display CNV score and correlation.

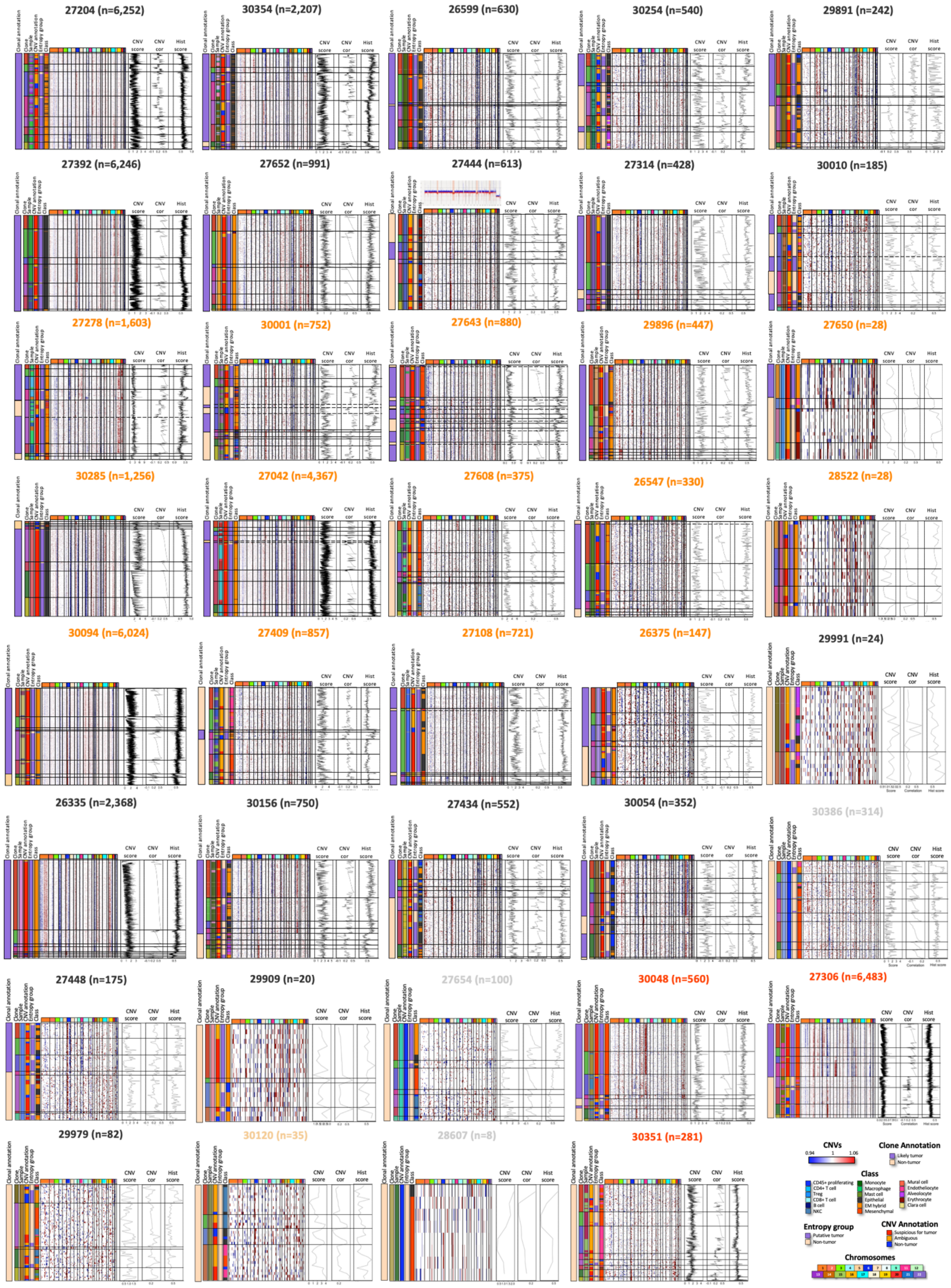

**Figure S5: Bulk CNV analysis confirms clonal CNV structure identified in scRNA-seq data using inferCNV, continued.** Each panel displays the inferCNV heatmap (bottom) and CNV profile obtained by Bionano analysis (top) for a MPM case. Heatmaps (cells x gene) display smoothed and centered gene expression computed by inferCNV, where genes are ordered by chromosomal position (indicated above). Cells are hierarchically clustered over a Euclidean metric with complete linkage and annotated with phenotype, cloneID, and tumor/non-tumor call. Line plots display the relative read density computed by Bionano software for the case subset to regions reported by inferCNV.

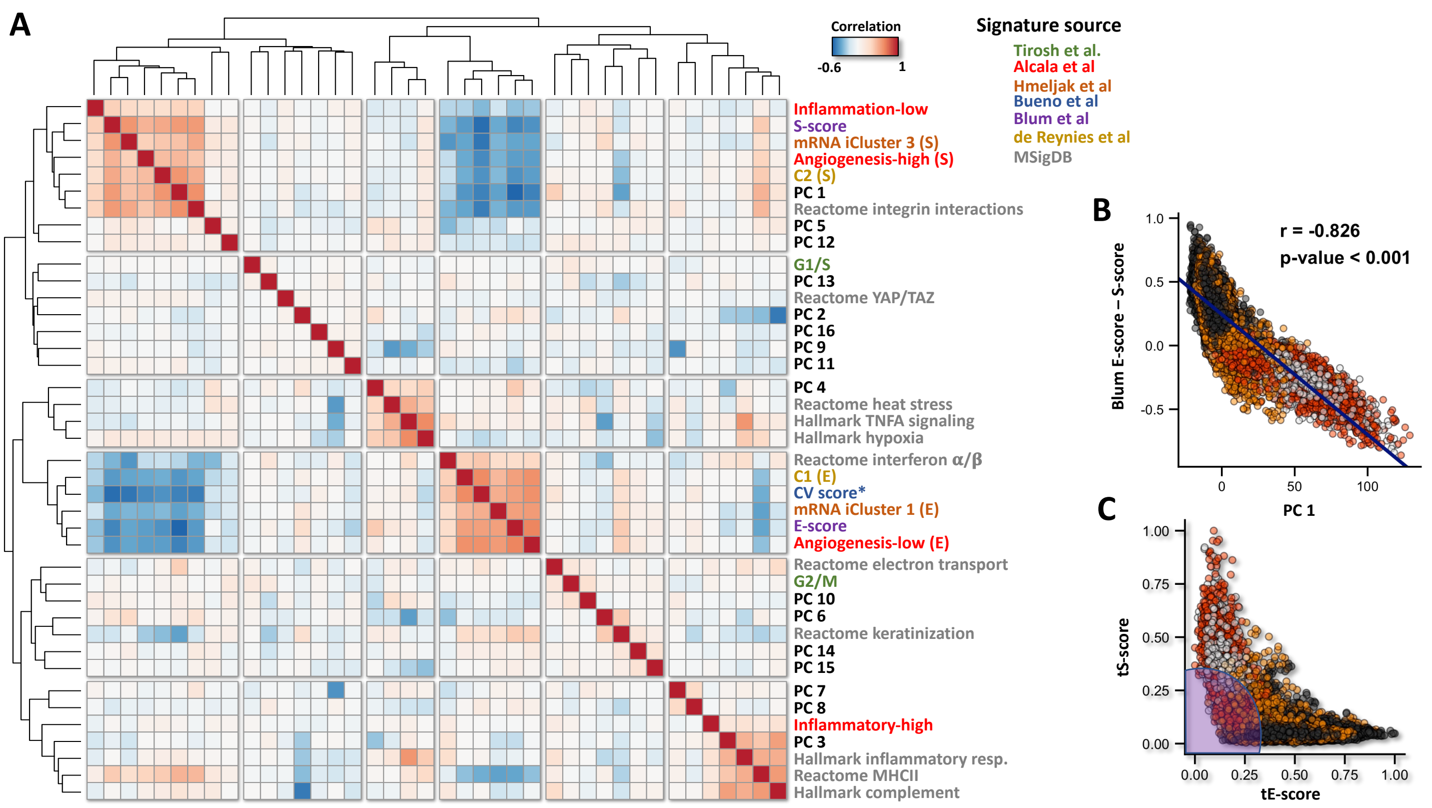


**Figure S6: Correlation of principal components and gene signatures across malignant cells, related to Figure 2. A)** Heatmap displays correlation between principal component embeddings, single cell scores for previously published signatures in MPM, and gene sets enriched in principal component loadings. Database or publication source of signatures are annotated by color. **B**) Relationship between Blum EM gradient score and PC1. **C)** Relationship between scRNA-seq derived sarcomatoid (tS) and epithelioid (tE) scores.

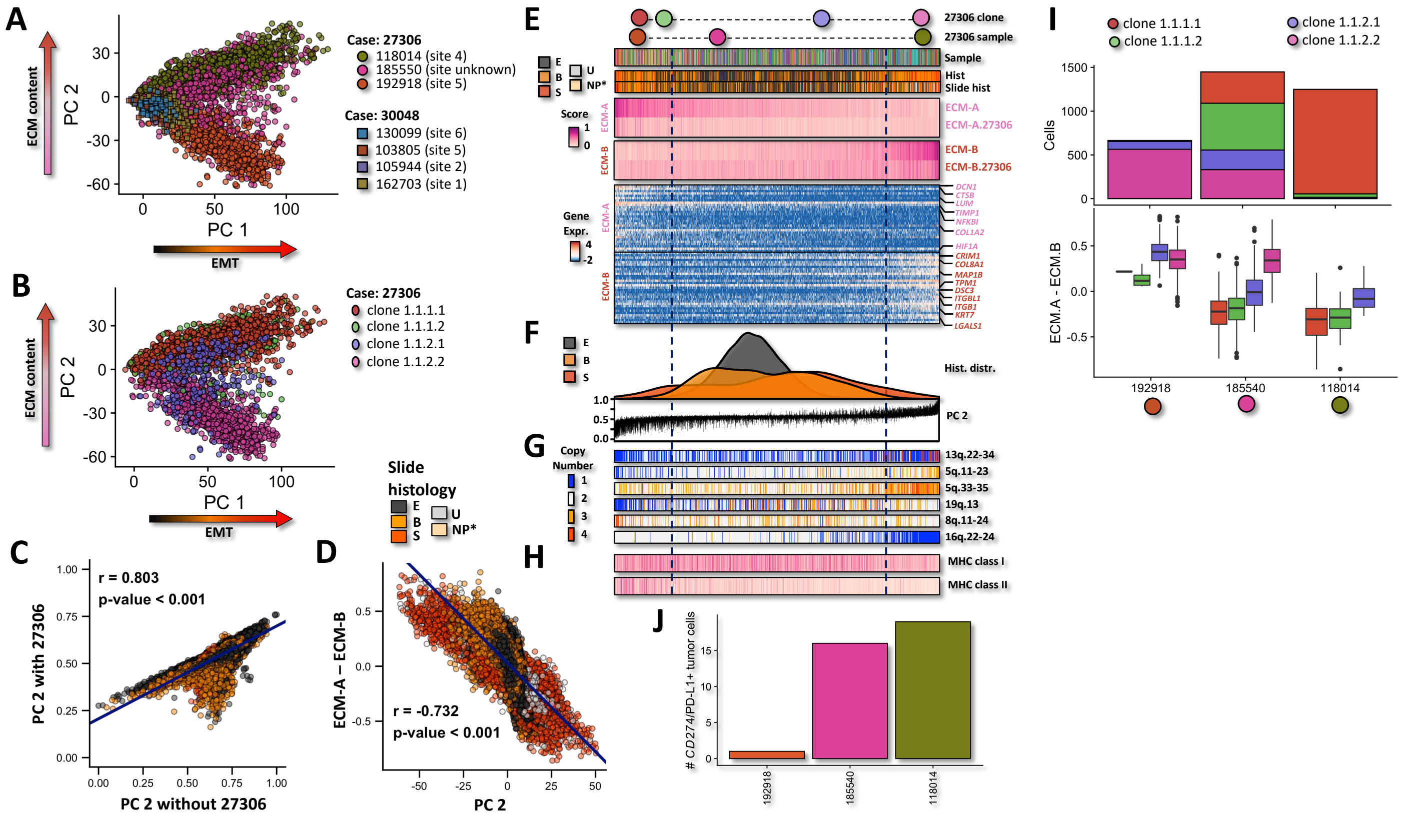

**Figure S7: ECM modules are variably expressed independent of histology and histomolecular EM gradient, related to Figure 2. A-B)** PCA of tumor cells isolated from sarcomatoid cases where either case and sample **(A)** or inferred CNV clone **(B)** are annotated. **C)** Correlation of second principal components where PCA was performed with and without 27306. **D)** Correlation between ECM score and second principal component. **E)** ECM signatures and associated gene expression, histology, and sample of origin. Above the relative positions of clones and samples of cells isolated from 27306 are indicated. Cells are ordered by ECM score and annotated with PC 2 scores and distribution of histology diagnosis **(F)** as well as relevant inferred CNVs **(G)**. **H)** Cell scores for KEGG MHC class I and II signatures are displayed. **I)** Clonal composition (bottom) and the EMC score distribution (top) of case 27306 cells by clone and sample. **J)** Number of CD274/PDL-1 positive tumor cells in each sample collected from 27306.

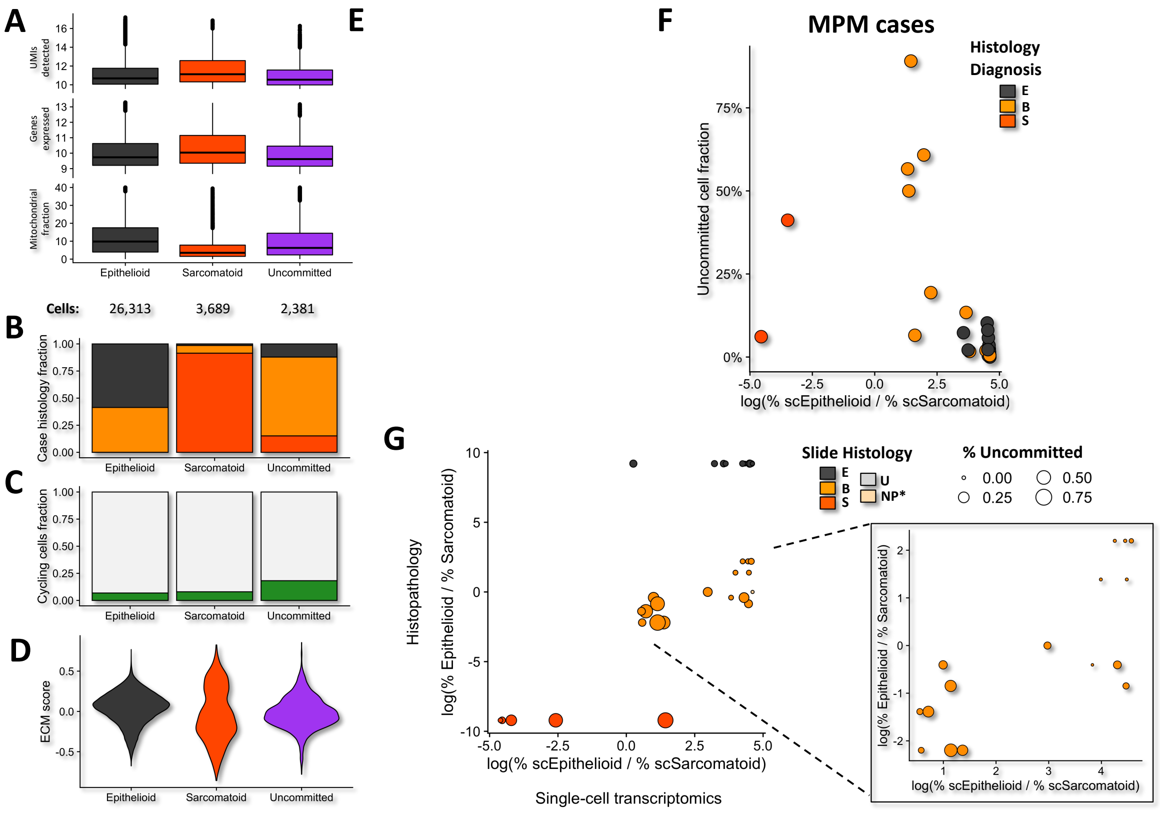


**Figure S8: A subset of MPM tumor cells are neither committed to sarcomatoid nor epithelioid programs, related to Figure 3. A)** Boxplots display the log2(UMIs detected), log2(genes expressed), and mitochondrial fraction observed in malignant cells of each malignant cell state. The number of cells in each state identified in cohort is listed below. **B)** Fraction of cells isolated from each PM histology diagnosis separated by malignant cell state (x-axis). **C)** Fraction of cells isolated with cycling signature by malignant cell state. **D)** Distribution of ECM score by malignant cell state. **E)** Candidate marker gene expression by cell type, where fraction of cell expressing gene and expression level are annotated by size and shade, respectively. **F)** Uncommitted cell fraction relationship with log ratio of fraction of Epithelioid and Sarcomatoid malignant cells in each case, where case diagnosis is annotated by color. **G)** Relationship between adjacent slide histopathologic analysis of log ratio of Sarcomatoid vs Epithelioid fraction with log ratio of fraction of Epithelioid and Sarcomatoid malignant cell by scRNA-seq in each case, where case diagnosis is annotated by color and uncommitted fraction is annotated by size.

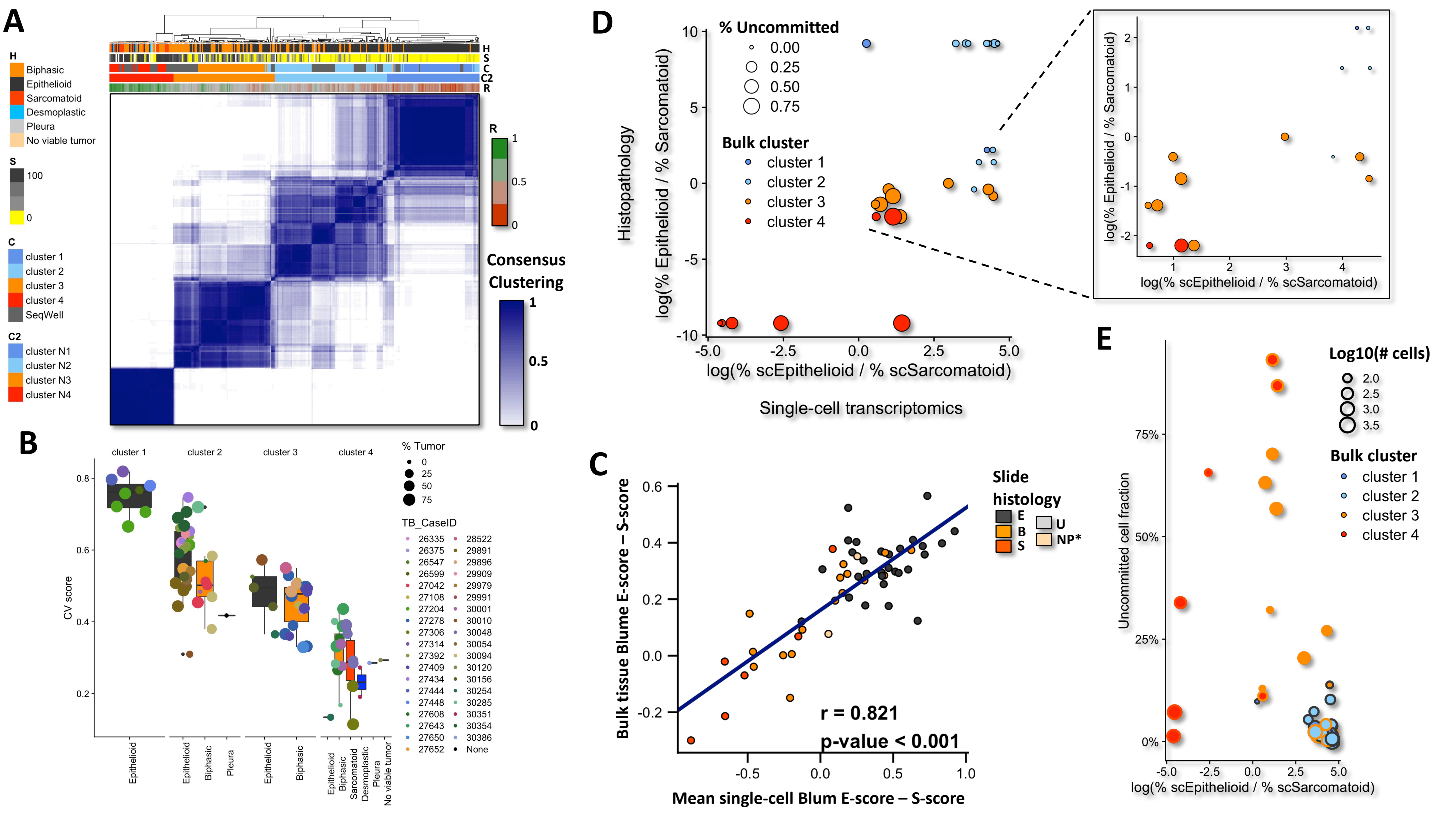

**Figure S9: Bulk RNA-seq assigns samples into EM-gradient and is concordant with scRNA-seq histomolecular profiles. A)** Consensus clustering of bulk RNA-seq samples with 211 previously published transcriptomes. Histology (H), % sarcomatoid (S), previously published transcriptional cluster assignment (C), current consensus cluster assignment (C2), and CV score (R) are annotated above the heatmap. **B**) Boxplot displays CV score and cluster assignment variation within cases. **C)** Correlation between bulk Blum EM gradient and mean Blum EM gradient of tumor single-cell transcriptomes for cases with greater than 30 tumor cells isolated. Dots are annotated by immediately adjacent slide histology. **D)** Correlation between E and S components as determined by scRNA-seq analysis of malignant cell fractions (x-axis) and histopathologic analysis of adjacent slides by PM sample. Samples are annotated with bulk transcriptional cluster (color) and fraction of uncommitted cells (size). Inset provides granular view of mixed malignant state samples. **E)** Uncommitted cell fraction relationship with log ratio of fraction of Epithelioid and Sarcomatoid malignant cells by scRNA-seq in each sample, where bulk transcriptional cluster (color and number of isolated cells per sample (size) are annotated.

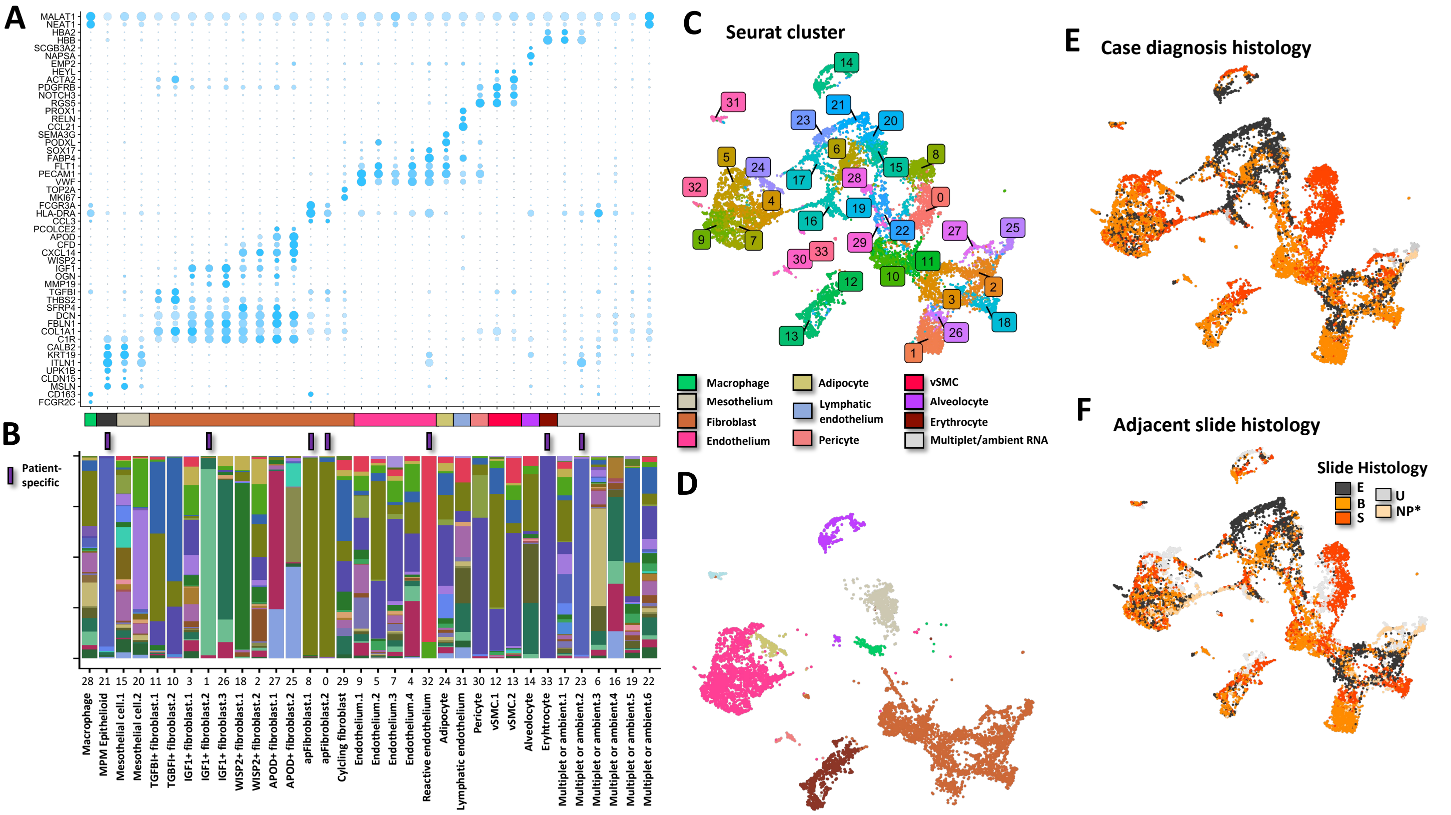


**Figure S10: Diverse stromal populations exist in PM tumors. A)** Select marker genes for MPM stromal clusters. Barplot below indicates stromal cell type for each cluster. **B**) Colors indicate fraction of cells within cluster isolated from each patient. Purple lines denote patient-specific clusters where 90% of cells within a cluster were isolated from a single patient. **C-F**) UMAPs display 12,776 high-quality stromal cells colored by Seurat cluster (**C**), stromal cell type (**D**), case histology diagnosis (**E**), or sample adjacent slide histology (**F**) in two dimensions. In **D**, cells from patient specific or multiplet/ambient RNA clusters are not visualized.

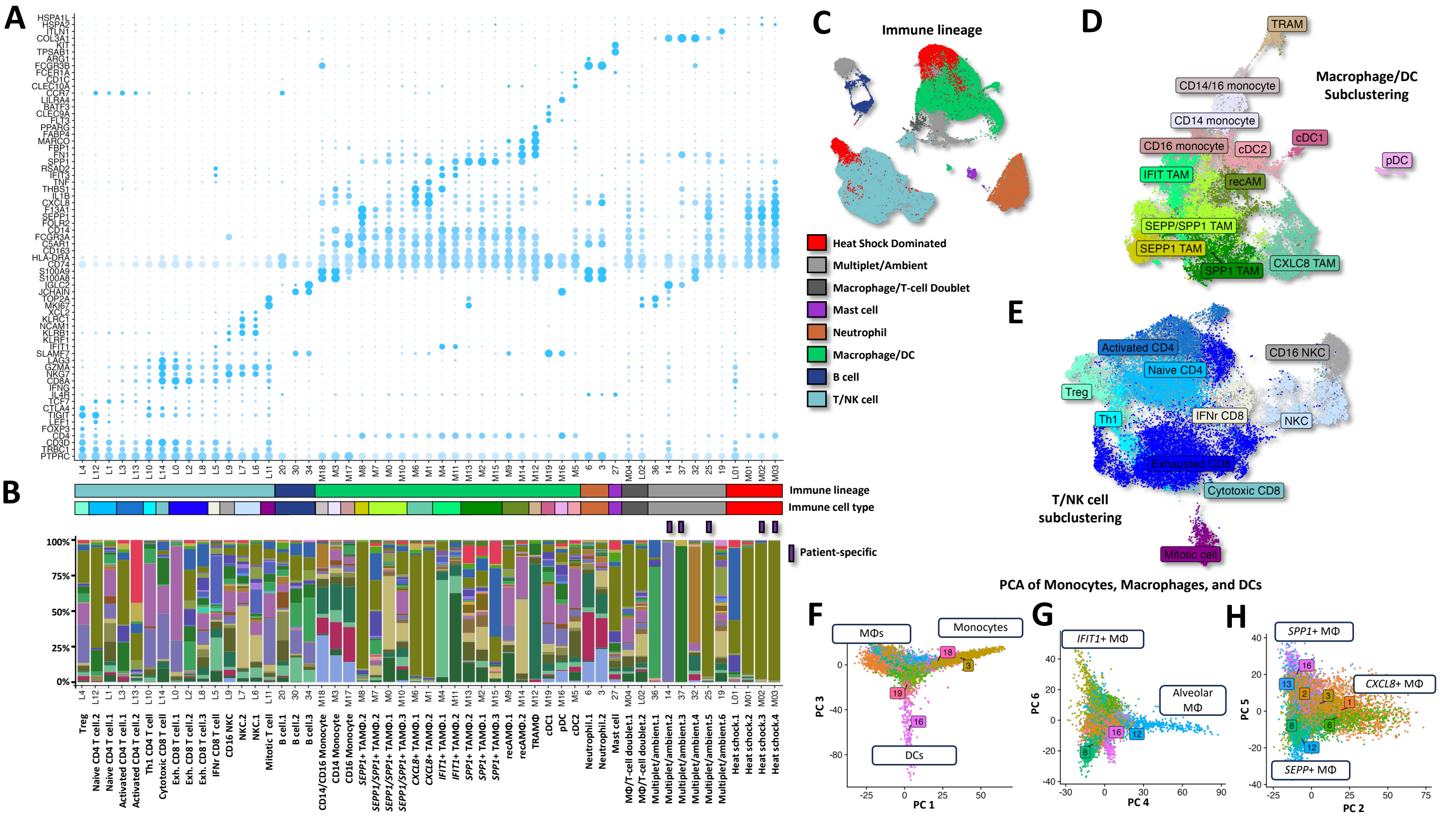


**Figure S11: Diverse immune populations exist in PM tumors. A)** Select marker genes for PM immune clusters and subclusters. Barplot below indicates the annotated broad immune lineage (above) and cell type (bottom) for each cluster. **B**) Colors indicate fraction of cells within cluster isolated from each patient. Purple lines denote patient-specific clusters where 90% of cells within a cluster were isolated from a single patient. **C**) UMAP displays 64,163 high-quality immune cells colored by annotated broad immune lineage in two dimensions. **D-E**) UMAPs display subclustered 16,922 monocytes, macrophages, and DCs **(D)** or 24,206 T- and NK cells **(E)** in two dimensions, where cells are colored by annotated cell type. **F-H**) PCA of monocytes, macrophages and DCs where scatterplots display cells plotted by principal component scores 1 vs 3 (**F**), 4 vs 6 (**G**), and 2 vs 5 (**H**). Cells are colored by graph-based Seurat myeloid, "M", subclusters.

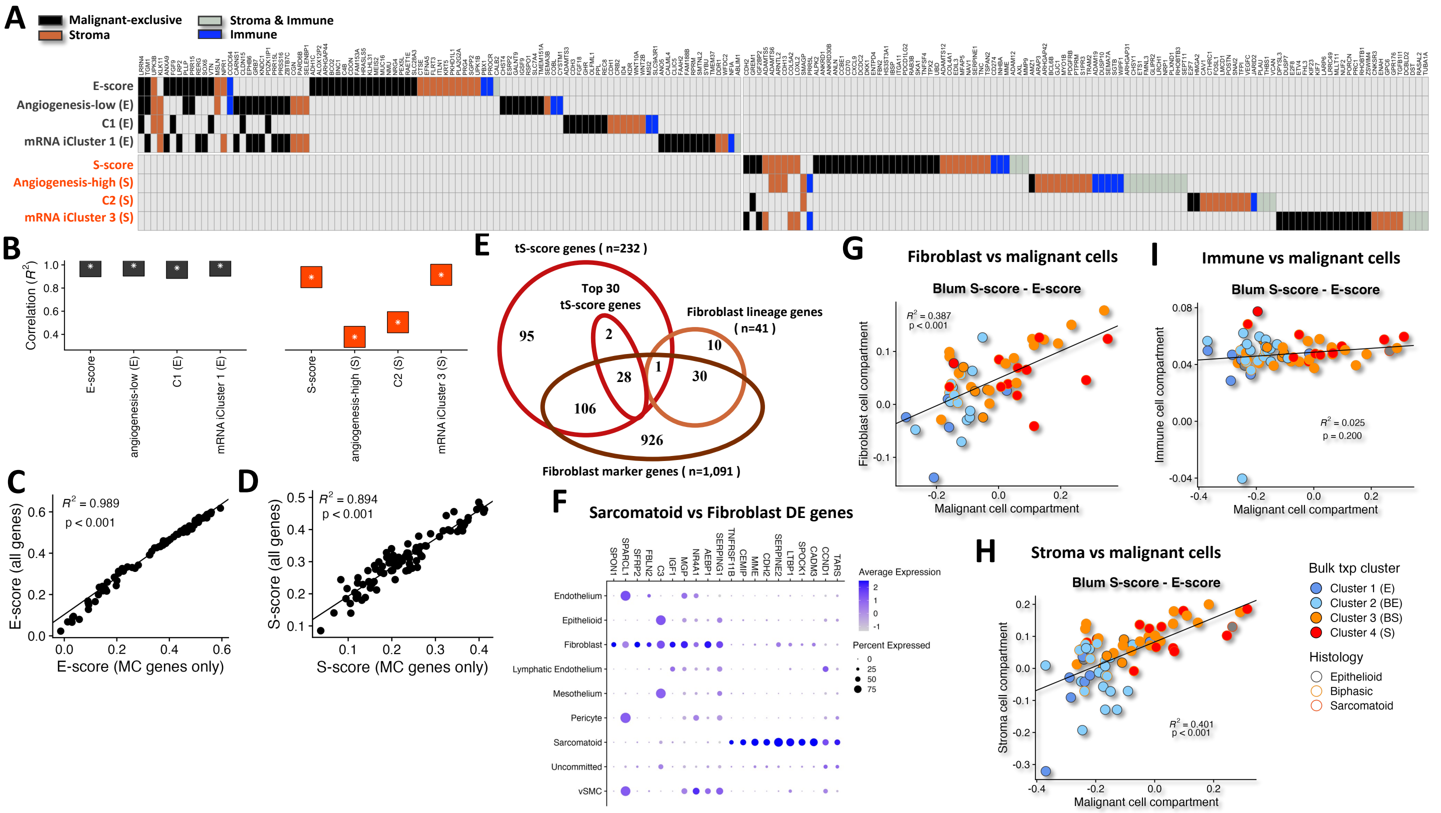


**Figure S12: Decomposition of bulk RNA-seq signatures and the tumor micro-environment. A)** Heatmap displays top 30 genes as columns from previously published EM histomolecular signatures (rows). Genes present in the EM histomolecular signature found in non-malignant components of the TME are colored: stroma (light brown), immune (blue), stroma & immune (light blue), exclusive to malignant cells (black). **B)** Boxes denote $R^2$ correlation values between EM histomolecular signatures and analogous signatures restricted only to malignant cell exclusive genes. **C-D)** Scatterplots display correlation between original and malignant exclusive E-score **(C)** and S-score **(D),** respectively. **E)** Venn diagram displays the overlap between malignant tS-score signature (all and top 30) with fibroblast marker and lineage genes identified in Seurat cluster marker and phenotype lineage analysis, respectively. **F)** Marker genes significantly differentially expressed (FC>1.5 & adj. p-value < 0.01) with more than 35% difference in fraction of cells expressing a given gene. **G-I)** Correlation between mean Blum S-score – E-score between either fibroblasts (**G**), entire stroma compartment (**H**), entire immune compartment (**I**), and malignant cells for each PM sample with sufficient cells of each type. Fill and outline colors reflect sample bulk transcriptional cluster and case histology, respectively.

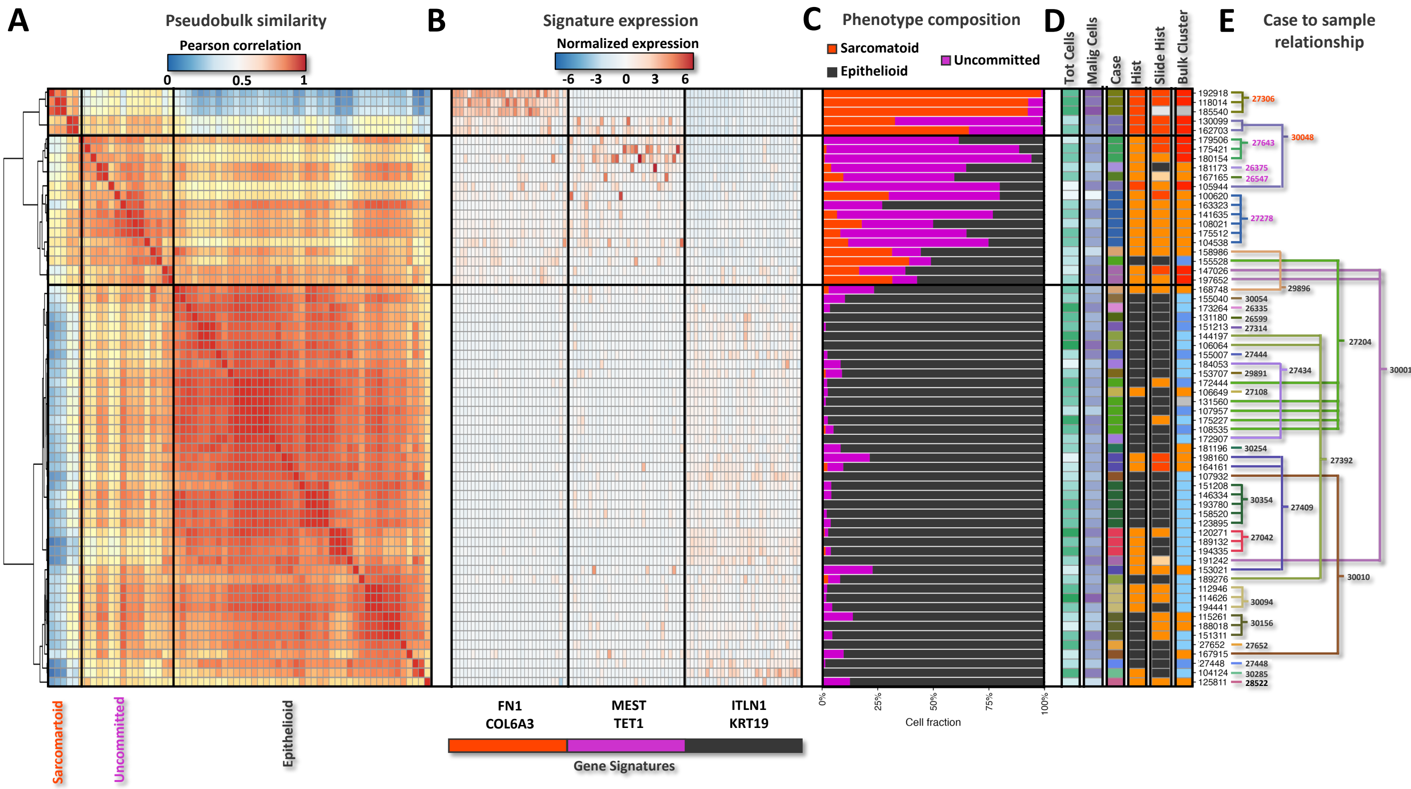


**Figure S13. PM tumor samples are categorized by average malignant cell state. A)** Hierarchically clustered correlation of mean aggregated expression of scRNA-seq malignant cell phenotype signatures for each PM sample (rows and columns). **B)** Mean aggregated expression of scRNA-seq malignant cell type signature genes (columns) for each PM sample (rows). **C)** Fraction of cells by scRNA-seq malignant cell phenotype within each PM sample (rows). **D)** Histology, PM case, bulk transcriptional and slide histology from adjacent tissue are annotated by color to the right**. E)** Tree diagram highlights samples isolated from the same surgical case.

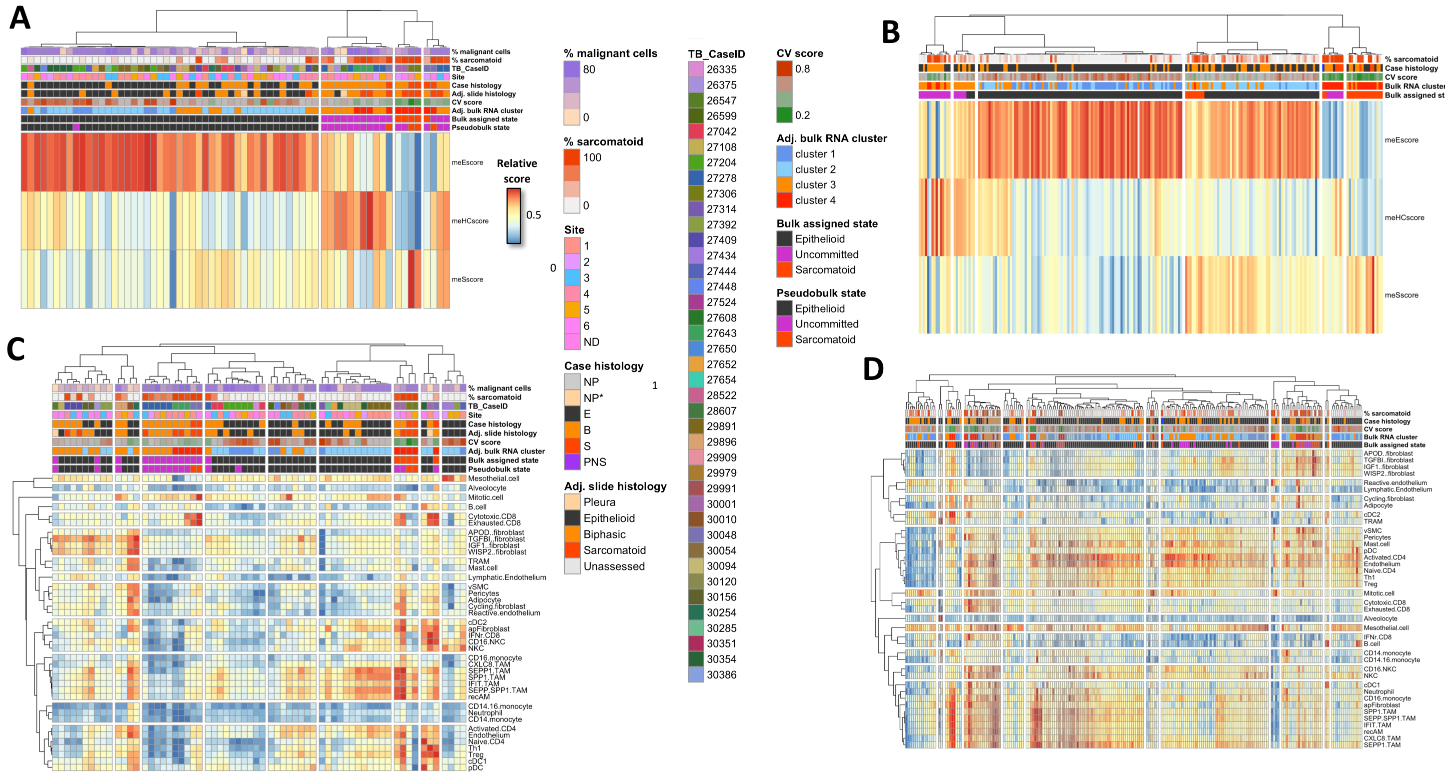


**Figure S14: scRNA-seq derived signatures deconvolute bulk RNA-seq samples. A-D)** Bulk RNA-seq samples from tissue adjacent to scRNA-seq samples (**A,C**) and 211 previously published PM samples (**B,D**) scored with scRNA-seq and previously published bulk RNA-seq signatures for malignant cell phenotypes (**A-B**) and cells within the TME (**C-D).**

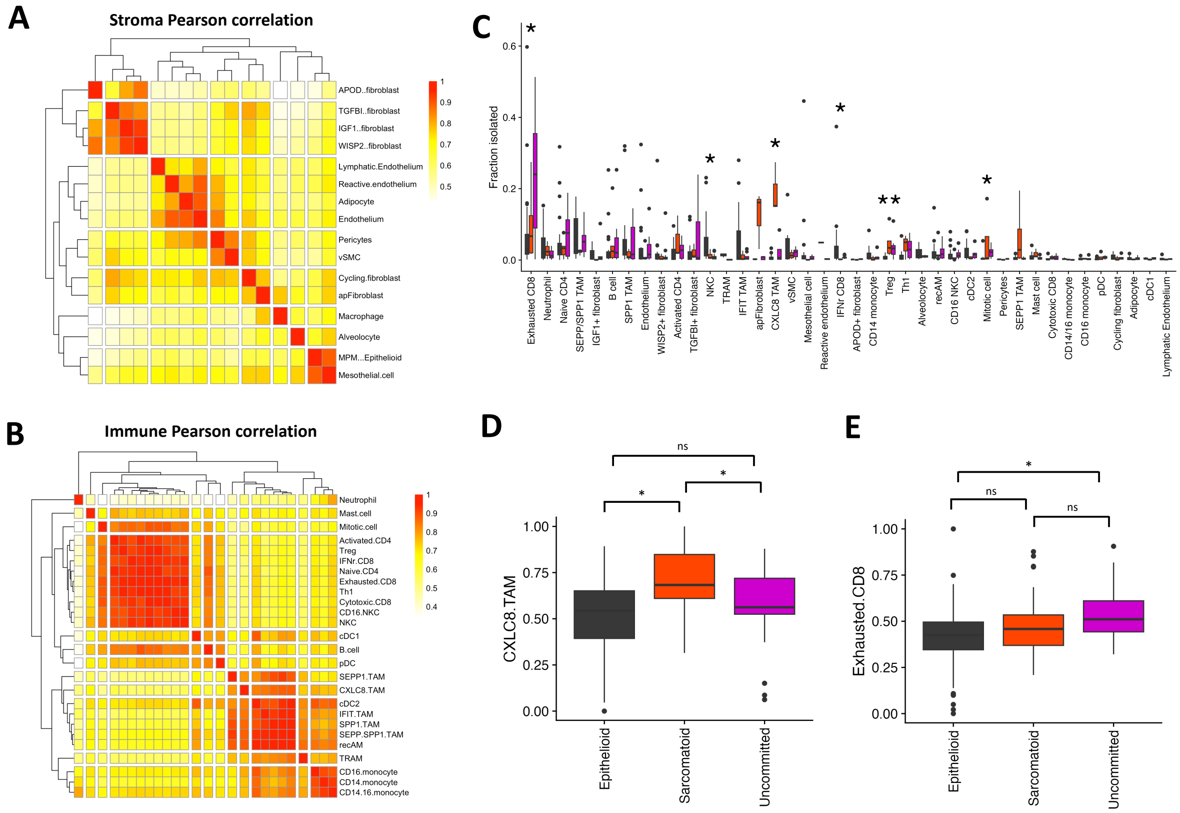


**Figure S15: PM TME varies with malignant cell state, related to Figure 4. A-B)** Pseudobulk representations of stromal (**A**) and immune (**B**) cell types are clustered by pearson correlation. **C**) Boxplots display fraction of cells isolated in the TME by scRNA-seq sample. **D-E**) Cell type signature scores of bulk RNA-seq samples from 211 previously published compared cross assigned bulk malignant cell state for CXCL8+ TAM (**D**) and exhausted CD8+ T cells (**E**). Asterisk indicates p-value <0.05 with Wilcoxon rank sum test.

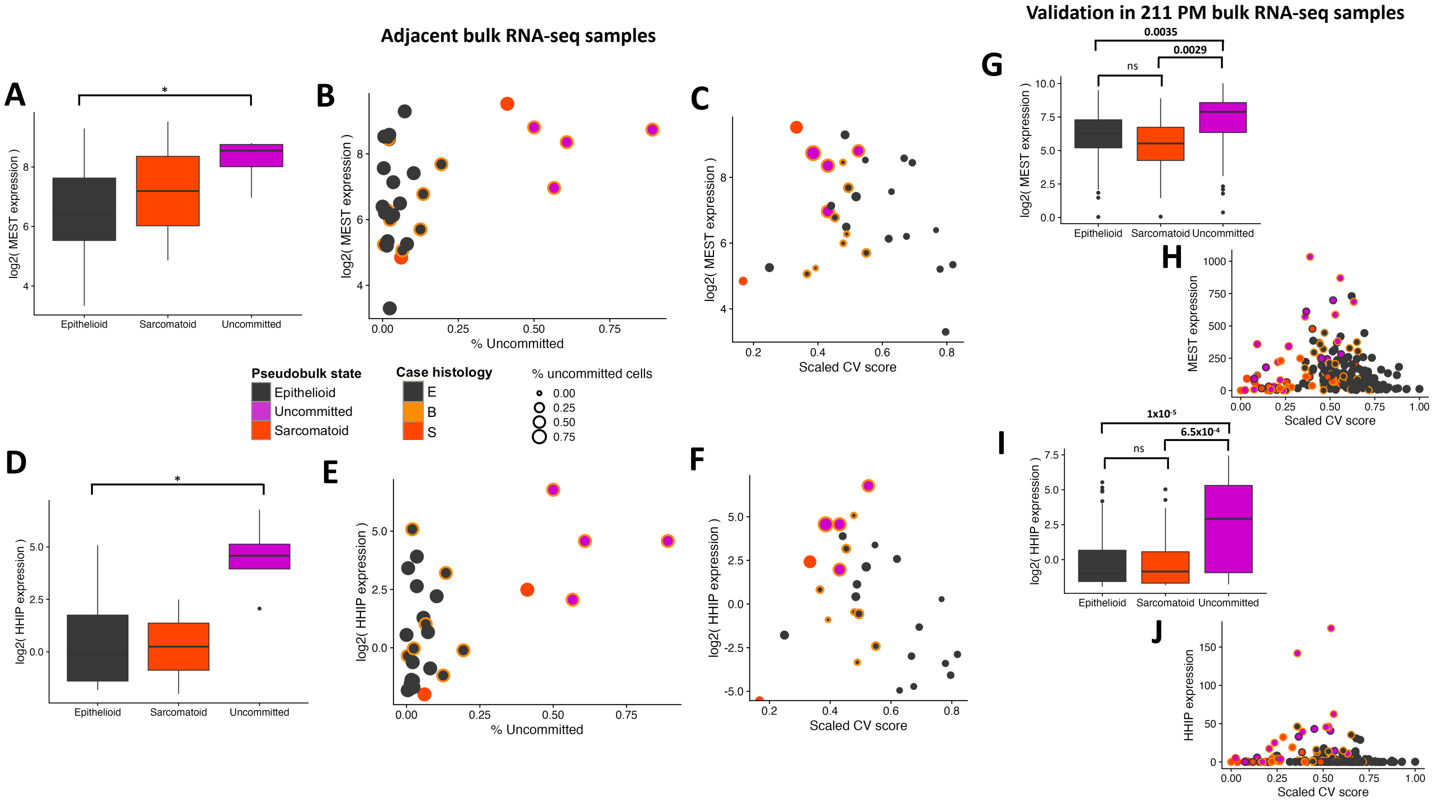


**Figure S16: *MEST* and *HHIP* operationalize uncommitted cells. A-F)** Biomarker assessment in adjacent bulk RNA-seq samples. **A,D)** Boxplots display log2 *MEST* (**A**) and *HHIP* (**D**) expression by predominant pseudobulk state. Asterisk denotes significance (p <0.05) with Wilcoxon rank sum test. **B-F**) Log2 *MEST* (**B-C**) and *HHIP* (**E-F**) compared to the fraction of uncommitted samples (**B,E**) or the scaled EM state as measured by Claudin-Vimentin (CV) ratio score (**C,F**). **G-J**) Validation of biomarkers in orthogonal cohort of 211 PM bulk RNA-seq cases. **G,I**) Boxplots display log2 *MEST* (**G**) and *HHIP* (**I**) expression by inferred predominant malignant cell state in independent bulk RNA-seq cohort. P-values are displayed for statistical comparison with Wilcoxon rank sum test. **H,J**) *MEST* (**H**) and *HHIP* (**J**) RNA expression 211 PM bulk RNA-seq cases vs scaled CV score. Fill and outline in scatterplots denote pseudobulk, actual or inferred, and case diagnosis, respectively.

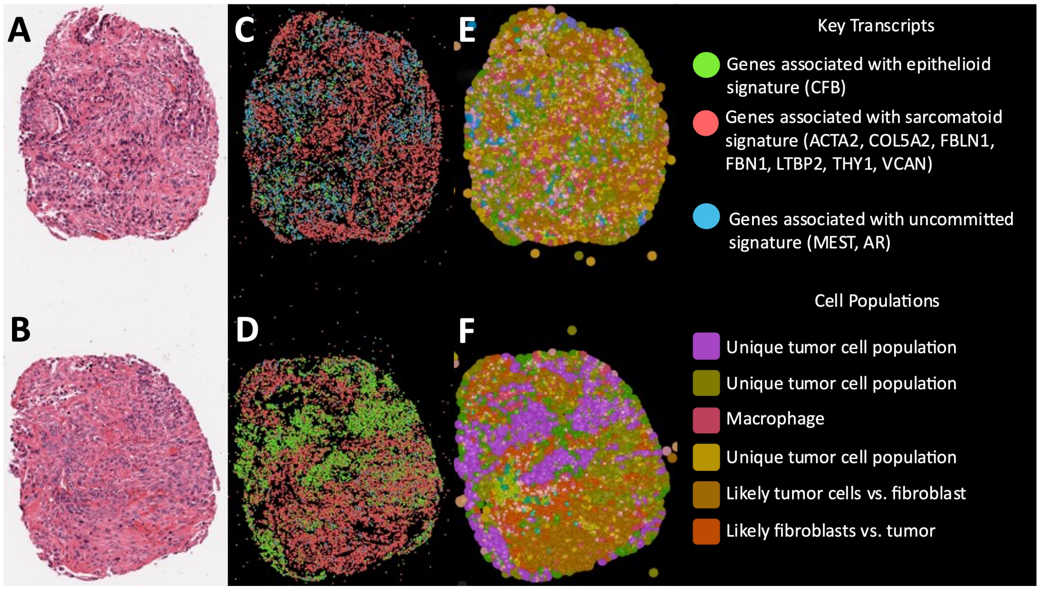


**Figure S17**: 10X Xenium Spatial transcriptomics of PM **A-B** H&E of two tissue cores from a single biphasic PM FFPE tumor block. **C-D** The expression of key transcripts derived from our single-cell study. **E-F** Clusters of unique cell populations. In these PM tumor cores, there exist multiple transcriptionally unique tumor cell populations with varying expression of MEST and other key transcripts.

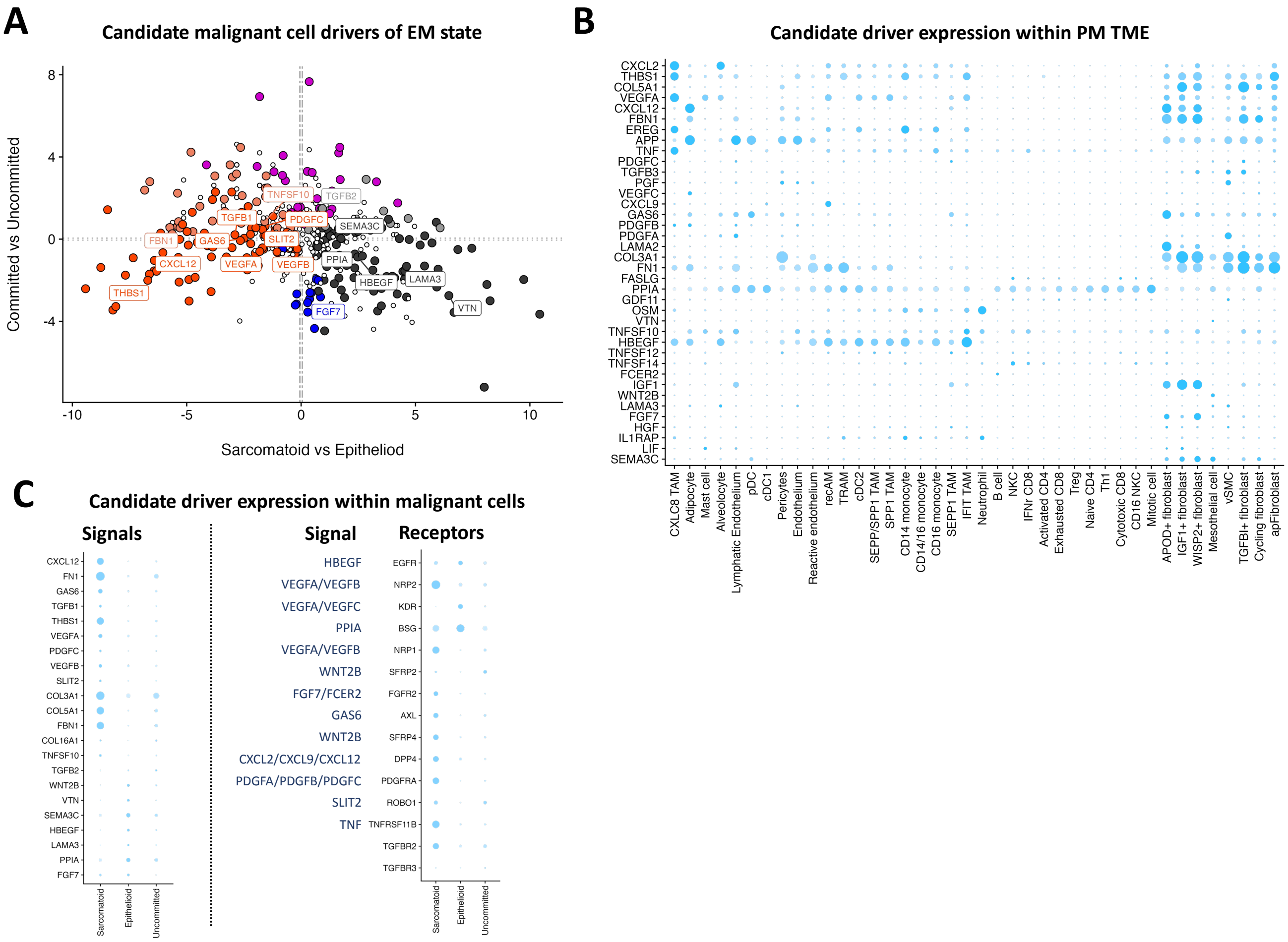


**Figure S18: Candidate drivers of malignant cell state. A)** TME cell type expression of signal proteins differentially expressed in non-malignant cells between tumors predominated by distinct malignant cell states. Labelled genes were identified as having significant TME interactions using CellPhoneDB. **B)** Log2 fold change gene expression of signal proteins in malignant cells between tumors of distinct pseudobulk malignant cell states. **C)** Malignant cell type expression of signal proteins differentially expressed in malignant cells between tumors predominated by distinct malignant cell states.
